# An unusual and physiologically vital protein with guanylate cyclase and P-type ATPase like domain in a pathogenic protist

**DOI:** 10.1101/475848

**Authors:** Özlem Günay-Esiyok, Ulrike Scheib, Matthias Noll, Nishith Gupta

## Abstract

Cyclic GMP is considered as one of the master regulators of diverse functions in eukaryotes; its architecture and functioning in protozoans remain poorly understood however. We characterized an unusual and extra-large guanylate cyclase (477-kDa) containing at least 4 putative P-type ATPase motifs and 21 transmembrane helices in a common parasitic protist, *Toxoplasma gondii*. This protein, termed as *Tg*ATPase_P_-GC due to its anticipated multi-functionality, localizes in the plasma membrane at the apical pole, while the corresponding cGMP-dependent protein kinase (*Tg*PKG) is distributed in cytomembranes. Both proteins are expressed constitutively during the entire lytic cycle of the parasite in human cells, which suggests a post-translational control of cGMP signaling. Homology modeling indicated an activation of guanylate cyclase by heterodimerization of its two cyclase domains. *Tg*ATPase_P_-GC is refractory to genetic deletion, and its CRISPR/Cas9-mediated disruption aborts the lytic cycle. Likewise, Cre/loxP-regulated knockdown of the *Tg*ATPase_P_-GC by 3’ UTR excision inhibited the parasite growth due to impairments in motility-dependent egress and invasion events. Consistently, cGMP-specific phosphodiesterase inhibitors restored the gliding motility of the mutant. A genetic repression of *Tg*PKG, or its pharmacological inhibition phenocopied the defects observed in the *Tg*ATPase_P_-GC mutant. Our data show a vital function of cGMP signaling, which is inducted by an alveolate-specific guanylate cyclase coupled to P-type like ATPase, and transduced by a dedicated PKG in *T. gondii*. The presence of *Tg*ATPase_P_-GC orthologs in many other alveolates with contrasting habitats implies a divergent functional repurposing of cGMP signaling in protozoans. The work also lays an avenue to systematically dissect the cascade and understand its evolution in a model protist.

## INTRODUCTION

Cyclic guanosine monophosphate (cGMP) is regarded as a common intracellular secondary messenger, which relays the endogenous and exogenous cues to the downstream mediators (kinases, ion channels *etc*.), and thereby regulates a range of cellular processes in prokaryotic and eukaryotic organisms (Hall and Lee 2018, Lucas et al. 2000). Initiating a physiological response to a biological cue requires a complex signal transduction beginning with the synthesis of cGMP from guanosine triphosphate (GTP) by the catalytic action of guanylate cyclase (GC). Levels of cGMP are strictly counterbalanced by phosphodiesterase enzyme (PDE), which degrades cGMP into GMP by hydrolyzing the 3’-phosphoester bond (Beavo 1995).

Much of our understanding of cGMP-induced transduction is derived from higher organisms, such as mammalian cells, which harbor four soluble guanylate cyclase subunits (α_1_, α_2_, β_1_, β_2_) and seven membrane-bound guanylate cyclases (GC-A to GC-G) (Lucas et al. 2000, Potter 2011). The soluble guanylate cyclase (sGC) isoforms function as heterodimers, which consist of an amino-terminal heme-binding regulatory domain, a dimerization region and a carboxyl-terminal catalytic domain. In contrast, all known particulate GC (pGC) proteins occur as homodimers except for GC-C, which exist as a homotrimer in the basal state (Lucas et al. 2000, Potter 2011). They usually have the following key features: an extracellular ligand binding site, a transmembrane domain, a kinase-homology regulatory sequence and a dimerization region, followed by a C-terminal catalytic domain comprising two functional catalytic sites. Nitric-oxide (NO) is the best-known activator of sGCs, while a variety of ligands are reported to activate pGCs (Lucas et al. 2000, Potter 2011). Protein kinase G (PKG or cGMP-dependent protein kinase) is a major actuator of cGMP signaling in most eukaryotic cells; it phosphorylates a repertoire of effector proteins to exert a consequent subcellular response (Lucas et al. 2000). All known PKGs belong to the serine/threonine kinases; in mammals, there are two variants (PKG I and PKG II). The type I PKGs, functioning as homodimers, have two soluble alternatively-spliced isoforms (α and β), encoded by a single gene (MacFarland 1995). The type II PKGs on the other hand are membrane-bound monomeric proteins encoded by different genes (MacFarland 1995, Pilz and Casteel 2003).

Unlike mammalian cells, little is understood about the overall architecture and functioning of cyclic nucleotide signaling in protozoans. The pathway seems to have diverged markedly in the course of evolution (Gould and de Koning 2011, Hopp et al. 2012, Linder et al. 1999). One of the protozoan phyla, Apicomplexa, which encompasses >6000 endoparasitic (mostly intracellular) species of significant clinical importance shows an even more intriguing design of cGMP signaling. *Toxoplasma*, *Plasmodium* and *Eimeria* are some of the key apicomplexan parasites, causing devastating diseases in humans as well as animals. These pathogens display a complex lifecycle in nature assuring their successful infection, reproduction, stage-conversion, adaptive persistence and inter-host transmission. Cyclic GMP cascade has been shown as one of the most central mechanism to coordinate the key steps during the parasitic lifecycle (Baker et al. 2017, Frénal et al. 2017, Gould and de Koning 2011, Govindasamy et al. 2016). In particular, the motile parasitic stages, *e.g*. sporozoite, merozoite, ookinete and tachyzoite deploy cGMP signaling to enter and/or exit host cells (Baker et al. 2017, Frénal et al. 2017), however the upstream actuation of the cascade is not yet understood.

Here, we identified an unusual guanylate cyclase linked to multiple P-type ATPase motifs in *T. gondii* and characterized its physiological importance for the asexual reproduction of the acutely infectious tachyzoite stage in mammalian host cells.

## MATERIALS AND METHODS

### Reagents and resources

The culture media and additives were purchased from PAN Biotech (Germany). Other common laboratory chemicals were supplied by Sigma-Aldrich (Germany). DNA-modifying enzymes were obtained from New England Biolabs (Germany). Commercial kits purchased from Life Technologies and Analytik Jena (Germany) were used for cloning into plasmids and for isolation of genomic DNA. Gene cloning and vector amplifications were performed in the XL1-Blue strain of *E. coli*. The RHΔ*ku80-hxgprt^−^* strain of *T. gondii*, lacking non-homologous end-joining repair (Fox et al. 2009, Huynh and Carruthers 2009) and thus facilitating homologous recombination-mediated genetic manipulation, was provided by Vern Carruthers (University of Michigan, USA). Fluorophore-conjugated secondary antibodies (Alexa488, Alexa594) and oligonucleotides were obtained from Thermo Fisher Scientific (Germany). The primary α-HA antibody and zaprinast were acquired from Sigma-Aldrich (Germany). cGMP-specific PDE inhibitor BIPPO (5-Benzyl-3-isopropyl-1H-pyrazolo [4,3*d*] pyrimidin-7(*6H*)-one) (Howard, Harvey, Stewart, Azevedo, Crabb, Jennings, Sanders, Manallack, Thompson and Tonkin 2015), and PKG inhibitor compound 2 (4-[7-[(dimethylamino) methyl]-2-(4-fluorophenyl) imidazo [1,2-α] pyridin-3-yl] pyrimidin-2-amine) (Biftu et al. 2005), were donated by Philip Thompson (Monash University, Australia) and Oliver Billker (Wellcome Trust Sanger Institute, UK), respectively. The primary antibodies against *Tg*Gap45 (Plattner et al. 2008) and *Tg*Sag1 (Dubremetz et al. 1993) were provided by Dominique Soldati-Favre (University of Geneva, Switzerland) and Jean-François Dubremetz (University of Montpellier, France), respectively. Human foreskin fibroblasts were obtained from Carsten Lüder (Georg-August University, Göttingen, Germany)

### Parasite and host cell cultures

Tachyzoites of *T. gondii* (RHΔ*ku80-hxgprt^−^* and its derivative strains) were serially maintained by infecting confluent monolayers of human foreskin fibroblasts (HFFs) every second day, as described previously (Gupta et al. 2005). Briefly, fresh host cells were harvested by trypsinization and grown to confluence in flasks, plates or dishes, as required for the experiments. Uninfected and parasitized HFF cells were cultivated in Dulbecco’s modified Eagle’s medium (DMEM) supplemented with 10% heat-inactivated fetal bovine serum (iFBS, PAN Biotech Germany), 2 mM glutamine, 1 mM sodium pyruvate, 1x minimum Eagle’s medium non-essential amino acids (100 µM each of serine, glycine, alanine, asparagine, aspartic acid, glutamate, proline), penicillin (100 U/ml), and streptomycin (100 µg/ml) in a humidified incubator (37 °C, 5% CO_2_). Freezer stocks of intracellularly replicating tachyzoites (HFFs with mature parasite vacuoles) were made in medium containing 45% iFBS, 5% DMSO and 50% DMEM including all supplements mentioned above.

### Genomic-tagging of *Tg*ATPase_P_-GC and *Tg*PKG

Sequences of *Tg*ATPase_P_-GC (TGGT1_254370) and *Tg*PKG (TGGT1_311360) genes were obtained from the parasite genome database (ToxoDB) (Gajria, Bahl, Brestelli, Dommer, Fischer, Gao, Heiges, Iodice, Kissinger, Mackey, Pinney, Roos, Stoeckert, Wang and Brunk 2008). The expression and subcellular localization of *Tg*ATPase_P_-GC and *Tg*PKG were determined by 3’-insertional tagging (3’IT) of corresponding genes with an epitope, essentially as reported before (Nitzsche et al. 2017). To achieve this, the 3’ end of the gene (1-1.5 kb 3’-crossover sequence or COS) was amplified from gDNA of the RHΔ*ku80-hxgprt^−^* strain using Q5^TM^ High-Fidelity DNA Polymerase (Bio-Rad Laboratories, Germany) (see Table S1 for primers). The HA-tagged amplicons were cloned into the *p3’IT-HXGPRT* vector using *Xcm*I/*EcoR*I and *Nco*I/*EcoR*I enzyme pairs for *Tg*ATPase_P_-GC and *Tg*PKG, respectively. This vector harbors 3’UTR of the *Tg*Gra1 gene (to stabilize the transcript) as well as a transgenic drug selection cassette expressing hypoxanthine-xanthine-guanine phosphoribosyltransferase (HXGPRT) as indicated in results section (Figure 2A, Figure S5A). The final constructs were linearized (15 µg) in the first-half of COS using *Sac*I (*Tg*ATPase_P_-GC) and *Sph*I (*Tg*PKG), followed by transfection into tachyzoites of RHΔ*ku80-hxgprt^−^* strain (10^7^). The fresh extracellular parasites were prepared by centrifugation (420*g*, 10 min) and suspended into sterile-filtered cytomix (120 mM KCl, 0.15 mM CaCl_2_, 10 mM K_2_HPO_4_/KH_2_PO_4_, 25 mM HEPES, 2 mM EGTA and 5 mM MgCl_2_, pH 7.4) supplemented with fresh ATP (2 µM) and glutathione (2 µM). They were transfected by electroporation using the BTX instrument (T-016 program, 2 kV, 50 Ω, 25 uF, 250 µs). Transgenic parasites expressing HXGPRT were selected with mycophenolic acid (*MPA, 25 µg/ml) and xanthine (50 µg/ml)* (Donald et al. 1996).

A successful epitope-tagging of the genes was verified by recombination-specific PCR and sequencing of subsequent amplicons. The stable drug-resistant transgenic parasites were subjected to limiting dilution to obtain the clonal lines for downstream analyses. The eventual strains expressed *Tg*ATPase_P_-GC-HA_3’IT_ or *Tg*PKG-HA_3’IT_ under the control of their native promoter and *Tg*Gra1-3’UTR. Using a similar strategy, we generated additional transgenic strains, in which *Tg*Gra1-3’UTR was replaced by the native 3’UTR of *Tg*ATPase_P_-GC and *Tg*PKG genes. Here, approximately 1 kb of 3’UTR beginning from the translation stop codon of the *Tg*ATPase_P_-GC and *Tg*PKG genes was amplified from gDNA of the RHΔ*ku80-hxgprt^−^* strain and then cloned into the *p3’IT-HXGPRT* plasmid at *EcoR*I/*Spe*I sites, substituting for *Tg*Gra1-3’UTR (primers in Table S1). The constructs were linearized and transfected into parasites, followed by transgenic selection, crossover-specific PCR screening, and clonal dilution, as described above.

### CRISPR/Cas9-assisted genetic disruption of *Tg*ATPase_P_-GC

The mutagenesis of *Tg*ATPase_P_-GC was achieved using CRISPR/Cas9 system, as reported previously for other genes (Sidik et al. 2014). To express gene-specific gRNA and Cas9, we utilized *pU6-sgRNA-Cas9* vector. The oligonucleotide pair, designed to target the nucleotide region from 145 to 164, was cloned into vector by golden gate assembly method using *Tg*ATPase_P_-GC-sgRNA-F1/R1 primers (Table S1). The assembly was initiated by mixing the *pU6-sgRNA-Cas9* vector (45 ng), *Bsa*I-HF enzyme (5 U) in CutSmart buffer (NEB), oligonucleotides (0.5 µM), T4 ligase (5 U) in ligation buffer (Thermo Fisher Scientific) in a total volume of 20 µl. The conditions were set as 30 cycles of 37°C – 2 min/ 20°C – 5 min followed by 37°C – 10 min/ 50°C – 10 min/ 80°C – 10 min. The product was directly transformed into XL1B strain of *E. coli*. Positive clones were verified by DNA sequencing, followed by transfection of 15 µg construct into P_native_-*Tg*ATPase_P_-GC-HA_3’IT_-*Tg*Gra1_3’UTR_ line (stated as the progenitor strain in Figure 3A) to disrupt the gene. A Cas9-mediated cleavage in the targeted locus caused a loss of HA-signal in transfected parasites, which was monitored by serial immunofluorescence assays at various time points of cultures, as indicated in the figure 3A.

### Cre-mediated knockdown of *Tg*ATPase_P_-GC and *Tg*PKG by excision of 3’UTR

The direct knockout of the *Tg*ATPase_P_-GC and *Tg*PKG genes led to mortality in tachyzoites (haploid genome), hence we performed knockdowns of the genes of interest (GOI) by Cre recombinase-mediated excision of the floxed 3’UTR in the HA-tagged strains. The progenitor strains expressing P_native_-*Tg*ATPase_P_-GC-HA_3’IT_-3’UTR_floxed_ or P_native_-*Tg*PKG-HA_3’IT_-3’UTR_floxed_ were transfected with a plasmid expressing Cre-recombinase, which recognizes and cuts off the loxP sites flanking 3’UTR and HXGPRT expression cassette (Figure 4A and Figure S5A). Tachyzoites transfected with Cre-encoding vector were then negatively selected for the loss of HXGPRT expression using 6-thioxanthine (80 µg/ml) (Donald, Carter, Ullman and Roos 1996). The single clones with Cre-excised 3’UTR were screened by PCR using indicated primers (Table S1) and validated by DNA sequencing. The reduction in the expression level of target proteins in the mutants was confirmed by immunofluorescence and immunoblot analysis, as described elsewhere.

### Measurement of cGMP in tachyzoites

Confluent HFF monolayers were infected either with the progenitor (P_native_-*Tg*ATPase_P_-GC-HA_3’IT_-3’UTR_floxed_; MOI: 1.5), or the knockdown mutant (P_native_-*Tg*ATPase_P_-GC-HA_3’IT_-3’UTR_excised_, MOI: 2) strain for 32-36 hours. The infected cells containing mature parasite vacuoles were then washed twice with ice-cold PBS, scraped by adding 2 ml colorless DMEM, and extruded through a 27G syringe (2x). Free tachyzoites were purified from the remaining host cell debris by low-speed centrifugation (420*g*, 10 min, 4°C). The parasite pellets were dissolved in 100 µl of chilled colorless DMEM for counting and cGMP extraction. The parasite suspension (5×10^6^, 100 µl) was mixed with 200 µl of ice-cold 0.1 M HCl and flash-frozen in liquid nitrogen. Samples were then thawed and squirted through pipette to disrupt the parasite membranes. Just the colorless DMEM and HFF cells, treated in the same way, were used as negative controls. Samples were transferred onto centrifugal filters (0.22-µm, Corning Costar Spin-X, CLS8169, Sigma) to eliminate the membrane particulates (20800*g*, 10 min, 4°C). The flow-through was filtered once more *via* 10-kDa filter units (Amicon Ultra-0.5 ml filters, Millipore) to obtain pure samples (20800*g*, 30 min, 4°C). Parasite extracts were then subjected to ELISA using the commercial ‘Direct cGMP ELISA kit’ (ADI-900-014, Enzo Life Sciences). The acetylated (2 hours) format of the assay was run for all samples including the standards and controls, as described by the manufacturer. The absorbance was measured at 405 nm, the data were adjusted for the dilution factor (1:3), and analyzed using the microplate analysis tool (www.myassays.com).

### Indirect immunofluorescence analysis (IFA)

IFA with extracellular and intracellular parasites was executed, as described elsewhere (Kong et al. 2018). Freshly egressed tachyzoites were incubated (20 min) on the BSA-coated (0.01%) coverslips for extracellular IFA. Intracellular parasites were stained with confluent monolayers of HFFs grown on coverslips (24-32 h post-infection). Samples were fixed with 4% paraformaldehyde (PFA) (15 min) followed by neutralization with 0.1 M glycine in PBS (5 min). For standard IFA, samples were permeabilized with 0.2% triton × 100/PBS (20 min), and nonspecific binding was blocked by 2% BSA in 0.2% triton × 100/PBS (20 min). Afterwards, parasites were stained with specified primary antibodies (rabbit/mouse α-HA, 1:3000; rabbit α-*Tg*Gap45, 1:8000; mouse α-*Tg*Sag1, 1:10000; mouse α-*Tg*ISP1, 1:2000) suspended in 2% BSA in 0.2% triton × 100/PBS (1 h). Cells were washed 3x with 0.2% triton × 100/PBS, followed by incubation with respective Alexa488-and 594-conjugated secondary antibodies (1 h) and 3x washing steps with PBS. Eventually, samples were mounted with Fluoromount G including DAPI for nuclei staining, and imaged with Zeiss Apotome microscope (Zeiss, Germany).

To resolve the C-terminal topology of *Tg*ATPase_P_-GC, fresh extracellular parasites were stained with rabbit α-HA (1:3000) and mouse α-*Tg*Sag1 (1:10000) or rabbit α-*Tg*Gap45 (1:8000) and α-*Tg*Sag1 (1:10000) antibody combinations before and after permeabilization as described elsewhere (Blume et al. 2009). Briefly, 5 × 10^4^ tachyzoites were released on BSA-coated coverslips and fixed with 4% PFA including 0.05% glutaraldehyde. Permeabilized coverslips were subjected to immunofluorescence assay as indicated above; however, all solutions were substituted to PBS for non-permeabilized staining as principle of the assay as shown in Figure 2B. To find the exact localization of *Tg*ATPase_P_-GC protein, the IMC was separated from the PM by treating extracellular parasites with α-toxin from *Clostridium septicum* (20 nM, 2 h) (List Biological Laboratories, US) followed by fixation on BSA-coated coverslips. In both cases, the standard secondary antibody staining procedure was performed afterwards, as described for immunofluorescence assay

### Lytic cycle assays

All assays were set up with fresh syringed-released parasites, essentially the same as reported earlier (Arroyo-Olarte et al. 2015). Parasitized cultures (MOI: 2; 40-44 h post-infection) were washed with standard culture medium, scraped, and extruded through a 27G syringe (2x). For plaque assays, HFF monolayers grown in 6-well plates were infected with tachyzoites (150 parasites/well) and incubated for 7 days without movement. Cultures were fixed with ice-cold methanol (−80°C, 10 min) and stained with crystal violet for 15 min followed by washing with PBS. The plaque sizes were measured by using ImageJ software (NIH, USA). To set up replication assays, host cells grown on coverslips placed in 24-well plates were infected with 3×10^4^ parasites (24 h or 40 h) before fixation, permeabilization, neutralization, blocking and immunostaining with α-*Tg*Gap45 and Alexa594 antibodies, as described in ‘Indirect immunofluorescence analysis’ section. The replication rates were assessed by enumerating parasites in their vacuoles. To measure the gliding motility, 4×10^5^ parasites, suspended in Hank’s balanced salt solution (HBSS) with or without drugs (BIPPO, 55 µM; zaprinast, 500 µM; compound 2, 2 µM), were incubated first to let them settle (15 min, RT) and glide (15 min, 37°C) on the BSA-coated (0.01%) coverslips. Samples were subjected to IFA using α-*Tg*Sag1 and Alexa488 antibodies after fixation, as mentioned above. Motile fractions were counted on the microscope, and trail lengths were quantified by the ImageJ software.

For invasion and egress, host cell monolayers cultured on glass coverslips were infected with MOI:10 for 1 h or MOI:1 for 40–64 h, respectively. For invasion BIPPO (55 µM), zaprinast (500 µM) and compound 2 (2 µM) treatments were done during the 1 h incubation time; however, the effects of the compounds on the parasite egress were tested 40 h post-infection by 5:30 min incubation after treatment. Cells were subsequently fixed with 4% PFA (15 min), neutralized by 0.1 M glycine/PBS (5 min), and then blocked in 3% BSA/PBS (30 min). Noninvasive parasites or egressed vacuoles were stained with α-*Tg*Sag1 antibody (mouse, 1:10000, 1 h) prior to detergent permeabilization. Cultures were washed 3x with PBS, permeabilized with 0.2% triton × 100/PBS (20 min) followed by staining with α-*Tg*Gap45 antibody (rabbit, 1:8000, 1 h) to visualize intracellular parasites. Samples were washed and stained finally with Alexa488 and Alexa594-conjugated antibodies (1:10000, 1 h). The fraction of invaded parasites was counted by immunostaining with α-*Tg*Gap45/Alexa594 (red), but not with α-*Tg*Sag1/Alexa488 (green). The percentage of lysed vacuole was scored directly by counting α-*Tg*Sag1/Alexa488 (green) and dual-stained parasites.

### Immunoblot analysis

Standard western blot was performed to determine the expression level of *Tg*PKG-HA_3’IT_, whereas the dot blot analysis was undertaken for *Tg*ATPase_P_-GC-HA_3’IT_. For the former assay, the protein samples prepared from extracellular parasites (2×10^7^) were separated by 8% SDS-PAGE (at 120 V) followed by semi-dry blotting onto a nitrocellulose membrane (85 mA/cm^2^) for 3h. The membrane was blocked with 5% skimmed milk solution prepared in TBS/0.2% Tween 20 (1 h by shaking at RT) and stained with rabbit α-HA (1:1000) and mouse α-*Tg*Rop2 (1:1000) antibodies. For the dot blot, samples equivalent to 10^7^ parasites were spotted directly on the nitrocellulose membrane. The membrane was blocked in 1% BSA/TBS-0.05% Tween 20 solution for 1h, followed by immunostaining with rabbit α-HA (1:1000) and/or rabbit α-*Tg*Gap45 (1:3000) antibodies diluted in the same buffer. Proteins were visualized by Li-COR imaging after staining with IRDye^®^ 680RD and IRDye^®^ 800CW (1:15000) antibodies.

### Expression of GC1 and GC2 of *Tg*ATPase_P_-GC in *Escherichia coli*

Heterologous expression of the GC1 and GC2 domains was performed in M15 and BTH101 strains of *E. coli*, for protein purification and functional complementation, respectively (see results). Starting with the first upstream start codon (ATG), the ORFs of GC1 (2850-3244 bp), GC2 (3934-4242 bp) and GC1+GC2 (2850-4242 bp) domains were amplified from the tachyzoite mRNA (RHΔ*ku80-hxgprt*^−^). The first-strand cDNA was generated from the total RNA using oligo-T primers of a commercial kit (Life Technologies) and used for ORF-specific PCR (primers in Table S1). The ORFs were cloned into the *pQE60* vector at the *Bgl*II restriction site resulting in a C-terminal 6xHis-tag. To purify the recombinant proteins, 5 ml culture of the M15 strains (grown overnight at 37°C) were diluted to an OD_600_ of 0.1 in 100 ml of Luria–Bertani (LB) medium containing 100 µg/ml ampicillin and 50 µg/ml kanamycin and incubated at 37 °C for 4-5 h until to an OD_600_ of 0.4-0.6. The cultures were then induced with 0.1 mM Isopropyl β-D-1-thiogalactopyranoside (IPTG) overnight at 25°C. The cell lysates were prepared under denaturing conditions, and proteins were purified using Ni-NTA column purification system according to the manufacturer’s protocol (Novex by Life Technologies). Briefly, cells were harvested (3000*g*, 20 min, 4°C) and resuspended in 8 ml lysis buffer (6 M GuHCl, 20 mM NaH_2_PO_4_, 500 mM NaCl, pH 7.8) by shaking for 10 min. They were disrupted by probe sonication (5 pulses, 30 sec each with intermittent cooling) and flash-frozen in liquid nitrogen followed by thawing at 37°C (3x). Intact cells were removed by pelleting at 3000*g* for 15 min. Lysate-containing supernatant was loaded on Ni-NTA column pretreated with binding buffer (8 M urea, 20 mM NaH_2_PO_4_, 500 mM NaCl, pH 7.8) followed by two washings by 4 ml washing buffer (8 M urea, 20 mM NaH_2_PO_4_, 500 mM NaCl) with pH 6.0 and pH 5.3, respectively. Proteins were eluted then in 5 ml elution buffer (20 mM NaH_2_PO_4_, 100 mM NaCl, 10 % Glycerol, pH 7.8). The eluate was concentrated using centrifugal filters with 30-kDa cut-off (Amicon ultra filters, Merck Millipore) by adding 4x volume of 20 mM NaH_2_PO_4_/100 mM NaCl/10% Glycerin to reduce urea concentration gradually in three centrifugation steps followed by storage at −80°C in liquid nitrogen. About 5 µg of purified protein was used for immunoblot analysis using mouse α-His (1:2000) (Dianova, Germany) and Alexa488 antibodies.

The function of GC1, GC2 and GC1+GC2 as being potential adenylate cyclase domains was tested in the *E. coli* BTH101 strain, which lacks cAMP signaling due to enzymatic deficiency and thus unable to utilize maltose as a carbon resource (Karimova, Pidoux, Ullmann and Ladant 1998). The *pQE60* constructs encoding indicated ORF sequences were transformed into the aforementioned bacterial strain. BTH101 cultures were grown overnight in 5 ml LB medium containing 100 µg/ml ampicillin and 100 µg/ml streptomycin at 37°C. Protein expression was induced by incubating cultures with 200 µM IPTG (2 h, 30°C) followed by dilution plating on MacConkey agar (pH.7.5) supplemented with 1% maltose, 200 µM IPTG and 100 µg/ml of each antibiotic. The strain harboring the empty *pQE60* vector served as a negative control, while the vector expressing *Cya*A (native bacterial adenylate cyclase) was included as a positive control. Agar plates were incubated at 30°C (~32 h) to examine for the appearing colonies.

### Bioinformatics and protein modeling

Protein sequences of *Tg*ATPase_P_-GC orthologs were obtained from the NCBI database (see figure S1 for gene IDs). A total of 30 guanylate cyclases were aligned and used for phylogenetic analysis based on the UPGMA clustering method by utilizing from the CLC Sequence Viewer 7.7 program. The tree was visualized then by Figtree v1.4.3. Similarly, sequence alignment of GC1 and GC2 domains of *Tg*ATPase_P_-GC was performed using Clustal Omega program (Sievers et al. 2011). The membrane topology of *Tg*ATPase_P_-GC was predicted by the combination of obtained data from TMHMM (Sonnhammer et al. 1998), SMART (Letunic et al. 2014), Phobius (Käll et al. 2004), NCBI conserved domain fsearch (Marchler-Bauer and Bryant 2004) and TMpred (Hofmann and Stoffel 1993) algorithms. Cyclase domain predictions for the homology model were based on Uniprot. GC1 (2929-3200 aa without loop standing 3038 and 3103) and GC2 (3989-4195 aa) domains were modeled using tmAC as the template (PDB code for tmAC. 1AZS; PDB code for ATPαS-bound tmAC, 1CJK; Q mean = −3.44; GMQE = 0.59; Swiss-Model). The ligand GTPαS was modeled into the pseudo-heterodimer.

### Data analyses and statistics

Graphs and underlying statistical significance were generated using GraphPad Prism v6.0. All experiments were performed at least three independent times, unless specified otherwise. Figures illustrating immunofluorescence images or making of the transgenic strains typically show a representative of three biological replicates. The error bars in graphs signify means with S.E.M. from multiple assays. The P-values were calculated by Student’s t-test, as reported in respective figure legends.

## RESULTS

### *T. gondii* encodes an alveolate-specific guanylate cyclase linked to a P-type ATPase

Our genome searches identified a single putative guanylate cyclase in the parasite database (ToxoDB) (Gajria et al. 2008), harboring a P-type ATPase like domain at its N-terminus and two nucleotide cyclase catalytic regions (designated as GC1 and GC2 based on the evidence herein, see below) at its C-terminus. Given the anticipated multi-functionality of this protein we named it *Tg*ATPase_P_-GC. The entire gene size is about 38.3-kb, consisting of 53 introns and 54 exons. The open reading frame (ORF) encodes for a remarkably large protein (4367 residues) with a predicted molecular weight of 477-kDa, comprising P-type ATPase (270-kDa) and nucleotide cyclase (207-kDa) domains, and includes a minimum of 21 transmembrane helices (Figure 1A). The first half of *Tg*ATPase_P_-GC (1-2480 aa) contains 10 α-helices and at least 4 conserved motifs: (a) The region from Lys^110^ to His^174^ encodes a phospholipid translocating ATPase; (b) the residues from Leu^207^ to Gly^496^ are predicted to form a bifunctional E1-E2 ATPase binding to both metal ions and ATP, and thus functioning like a cation-ATPase; (c) the amino acids from Thr^1647^ to Ser^1748^ harbor yet another metal-cation transporter with an ATP-binding region; (d) the motif from Cys^2029^ to Asn^2480^ comprises a haloacid dehalogenase-like (HAD-like) hydrolase, or a second phospholipid translocating ATPase.

**Figure 1.**
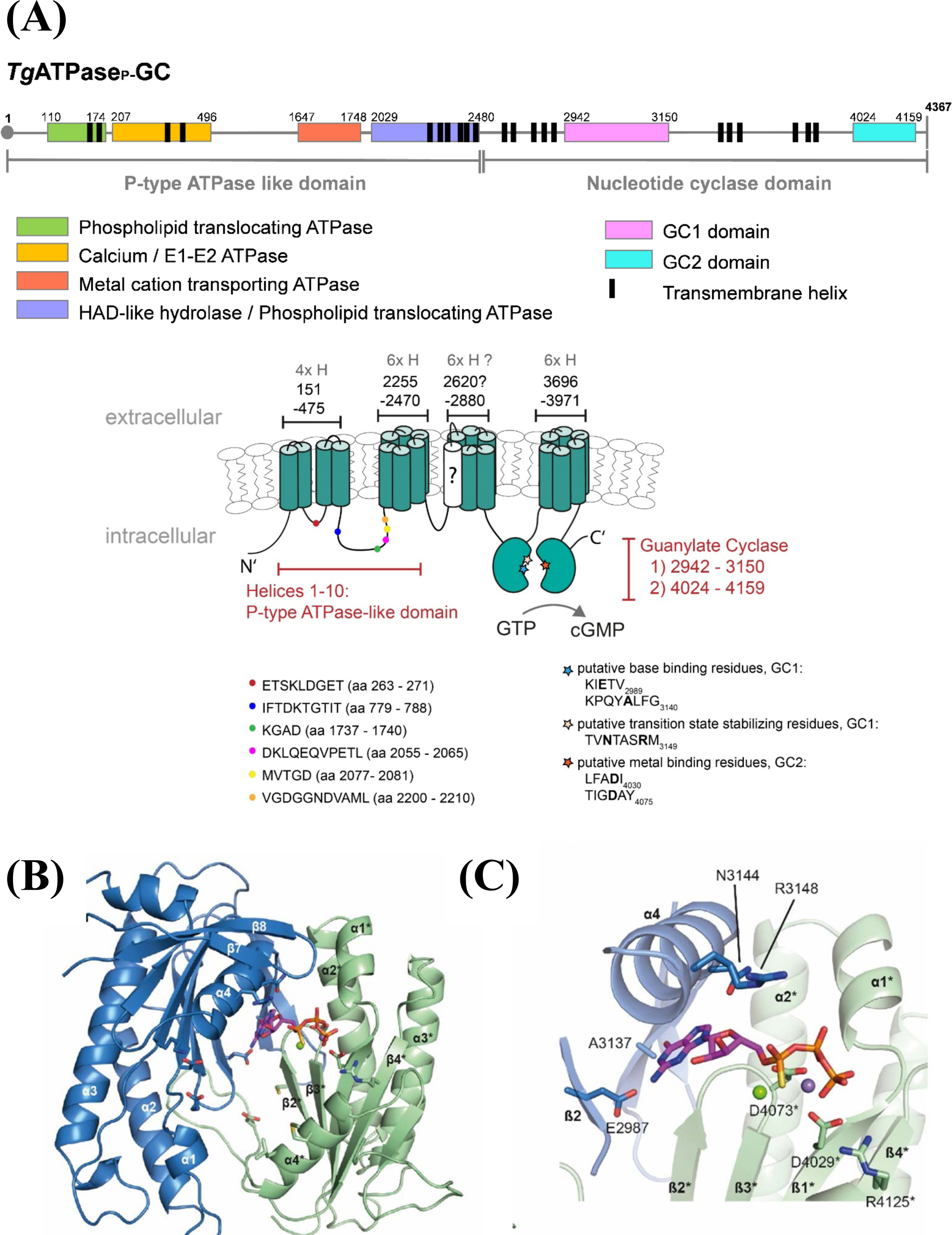
The genome of *Toxoplasma gondii* harbors an unusual heterodimeric guanylate cyclase conjugated to a P-type ATPase-like domain with multiple motifs. **(A)** The primary and secondary topology of *Tg*ATPase_P_-GC were predicted using TMHMM, SMART, TMpred, Phobius and NCBI domain search tools. The models were constructed by consensus across algorithms regarding the position of domains and transmembrane spans. The N-terminal half of the protein (1-2480 aa) containing 10 α-helices resembles a P-type ATPase with at least four motifs, as indicated by different colors. The C-terminal half (2481-4367 aa) harbors two potential nucleotide cyclase catalytic regions, termed GC1 and GC2, each with 6 transmembrane helices. Low-probability helix is question-marked. The color-coded signs on secondary structure show the position of highly conserved sequences in ATPases and cyclase domains. The key residues involved in base binding and catalysis of cyclases are also depicted in bold letters. **(B-C)** Tertiary structure of GC1 and GC2 domains based on homology modeling. The ribbon diagrams of GC1 and GC2 suggest a functional activation of the enzyme by pseudo-heterodimerization (similar to transmembrane adenylate cyclases (tmACs)). The model shows an antiparallel arrangement of GC1 and GC2, where each domain harbors a 7-stranded β-sheet surrounded by 3 α-helices. The image in *panel c* shows a GC1-GC2 heterodimer interface bound to GTPαS. Residues of GC2 labeled with asterisk (*) interact with the phosphate backbone of the nucleotide.

It is worth noting that a couple of single mutations were detected in the conserved ATPase sequences. The second motif of ATPase domain, which is predicted to be a calcium ATPase (yellow colored in figure 1A), showed two altered residues (ETS**K**L**D**GET) instead of ETSLLNGET. The beginning of subsequent cation transporting motif (highlighted with orange) contained alanine (Ala^1739^) in lieu of serine. Besides, one amino acid replacement (leucine to Ile^787^) was observed in the IFTDKTGTIT motif located in the linkage region of aforementioned motifs. It is also known that ATPase domains have an intrinsic kinase activity to phosphorylate itself on the aspartate residue in the ATP-binding pocket (Bublitz et al. 2011), which was found to be substituted to glutamate in the conserved DKLQ**E**QVPETL sequence located in the second predicted phospholipid-translocating ATPase motif (marked with blue in Figure 1A).

The second half (2481-4367 aa) includes a potential nucleotide cyclase comprising GC1 and GC2 domains from Ser^2942^-Lys^3150^ and Thr^4024^-Glu^4159^ residues, respectively (Figure 1A). Both GC1 and GC2 follow a transmembrane region, each with six helices. The question-marked helix antecedent to GC1 (2620-2638 aa) has a low probability. An exclusion of this helix from the model however results in a reversal of GC1 and GC2 topology (facing outside the parasite), which is unlikely given the intracellular transduction of cGMP signaling *via Tg*PKG. Moreover, our experiments suggest that C-terminal of *Tg*ATPase_P_-GC faces inwards (see figure 2B),. Phylogenetic study indicated an evident clading of *Tg*ATPase_P_-GC with homologs from parasitic (*Hammondia, Eimeria*, *Plasmodium*) and free-living (*Tetrahymena*, *Paramecium*, *Oxytricha*) alveolates (Figure S1). In contrast, the GCs from animals (soluble and receptor-type) and plants formed their own distinct clusters. Intriguingly, the protist clade contained two groups, one each for apicomplexans and ciliates, implying a phylum-specific evolution of GC proteins.

**Figure 2.**
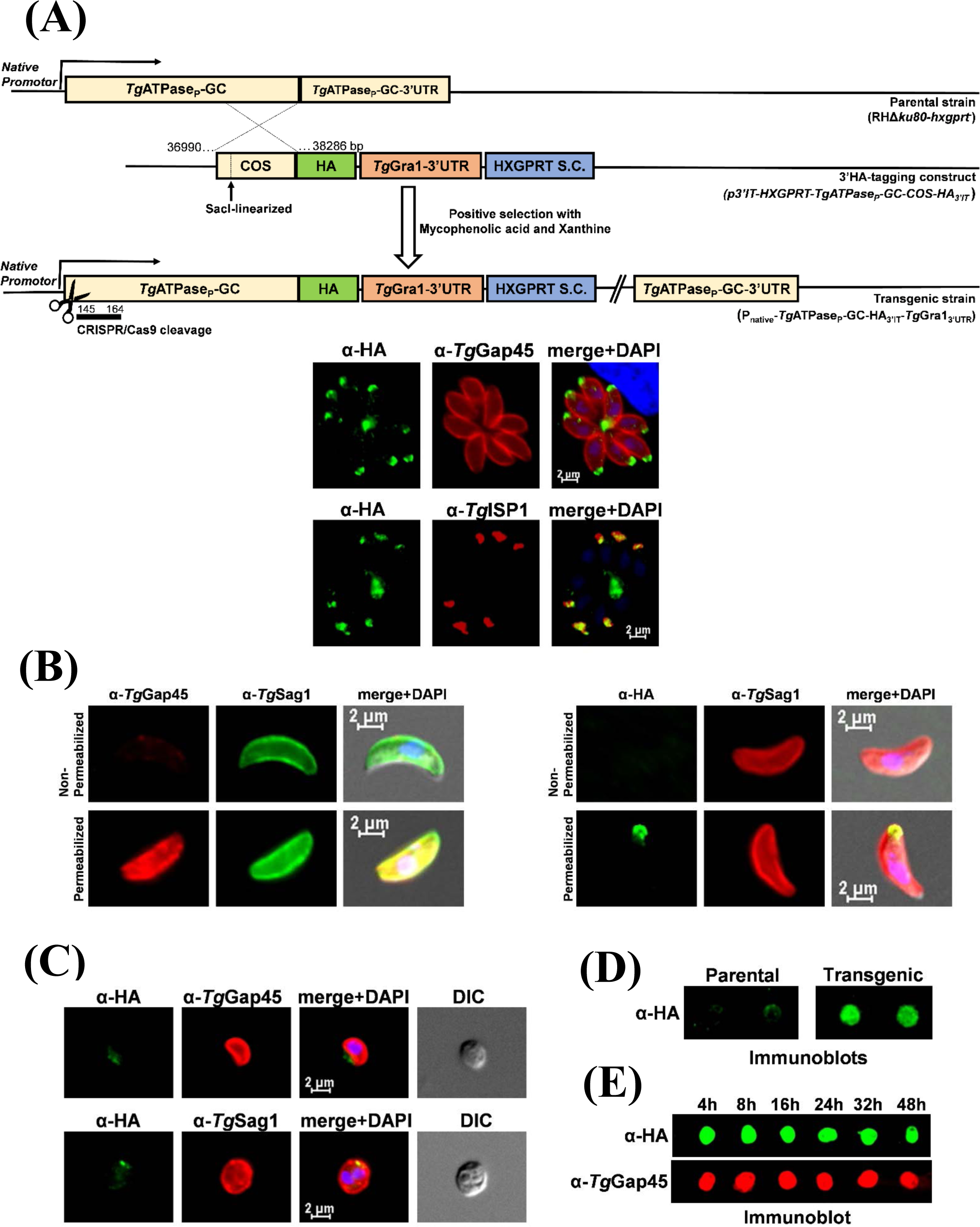
*Tg*ATPase_P_-GC is a constitutively expressed protein located in the plasma membrane at the apical pole of *T. gondii*. **(A)** Scheme for the genomic tagging of *Tg*ATPase_P_-GC with a 3’-end HA epitope. The *Sac*I-linearized plasmid for 3’-insertional tagging (*p3’IT-HXGPRT-TgATPase_P_-GC-COS-HA_3’IT_*) was transfected into parental (RHΔ*ku80-hxgprt^−^*) strain followed by drug selection. Intracellular parasites of the resulting transgenic strain (P_native_-*Tg*ATPase_P_-GC-HA_3’IT_-*Tg*Gra1_3’UTR_) were subjected to immunostaining with indicated antibodies (24 h post-infection). The host and parasite nuclei were stained by DAPI. *COS*, crossover sequence; *S.C*., selection cassette. **(B)** Immunofluorescence staining of extracellular parasites expressing *Tg*ATPase_P_-GC-HA_3’IT_ prior to and after detergent permeabilization. The α-HA staining of free parasites was performed before or after membrane permeabilization either using PBS without any BSA inclusion, or with 2% BSA dissolved in 0.2% triton × 100-supplemented PBS, respectively. No HA-staining was seen prior to permeabilization, which became apparent only after detergent treatment, confirming an inward orientation of the C-terminal of *Tg*ATPase_P_-GC (as predicted in figure 1A) **(C)** Immunostaining of extracellular parasites encoding *Tg*ATPase_P_-GC-HA_3’IT_ after drug-induced splitting of the inner membrane complex (IMC) from the plasma membrane (PM). Tachyzoites were incubated with α-toxin (20 nM, 2 h) prior to immunostaining with α-HA antibody in combination with primary antibodies recognizing IMC (α-*Tg*Gap45) or PM (α-*Tg*Sag1), respectively. **(D-E)** Immunoblots of tachyzoites expressing *Tg*ATPase_P_-GC-HA_3’IT_, and of the parental strain (RHΔ*ku80-hxgprt^−^*, negative control). The protein samples prepared from fresh extracellular tachyzoites (10^7^) were directly loaded onto the nitrocellulose membrane following staining with α-HA antibody. Samples shown in *panel e* were collected at different time points during the lytic cycle. α*-Tg*Gap45 antibody was used as loading control.

### GC1 and GC2 domains of *Tg*ATPase_P_-GC form a pseudo-heterodimer

The arrangement and architecture of GC1 and GC2 domains in *Tg*ATPase_P_-GC corresponds to mammalian transmembrane adenylate cyclase (tmAC) of the class III. The latter is activated by G-proteins to produce cAMP following extracellular stimuli (*e.g*. hormones). The cyclase domains of tmACs, C1 and C2, form an antiparallel pseudo-heterodimer with one active and one degenerated site at the dimer interface (Linder and Schultz 2003). Amino acids from both domains contribute to the binding site, and seven conserved residues play essential roles for nucleotide binding and catalysis (Linder and Schultz 2003, Sinha and Sprang 2006, Steegborn 2014). These include two aspartate residues, which bind two divalent metal cofactors (Mg^+2^, Mn^+2^) crucial for substrate placement and turnover. An arginine and asparagine stabilize the transition state, while yet another arginine binds the terminal phosphate (Pγ) of the nucleotide. A lysine/aspartate pair underlies the selection of ATP over GTP as the substrate. By contrast, a glutamate/cysteine or glutamate/alanine pair defines the substrate specificity in guanylate cyclases. The nucleotide binding and transition state stabilization are conferred by one domain, while the other domain directly or *via* bound metal ions interacts with the phosphates of the nucleotide in tmACs (Linder 2005, Sinha and Sprang 2006, Steegborn 2014).

The sequence alignment of GC1 and GC2 domains from *Tg*ATPase_P_-GC to their orthologous GCs/ACs showed that unlike other cyclases GC1 contains a 74-residues long loop insertion (3033-3107 aa) (Figure S2). The tertiary model structure (depleted for the loop inserted between α3 and β4 of GC1) shows that both domains consist of a seven stranded β-sheet surrounded by three helices (Figure 1B-C). The key functional residues with some notable substitutions could be identified as distributed in GC1 and GC2. In GC1 domain, one of the two metal binding (Me) aspartates is replaced by glutamate (E2991), while both are conserved in GC2 (D4029 and D4073) (Figure 1C and Figure S2). The transition state stabilizing (Tr) asparagine (N3144) and arginine (R3148) residues are located within GC1 domain (Figure 1C), however both are replaced by L4153 and M4157, respectively in GC2 motif (Figure S2). Another arginine (R4125) that is responsible for phosphate binding (Pγ) in tmACs is conserved in the GC2 domain (Figure 1C), whereas it is substituted by K3116 in GC1 (Figure S2). The cyclase specificity defining residues (B) are glutamate/alanine (E2987/A3137) and cysteine/aspartate (C4069/D4146) pairs in GC1 and GC2, respectively (Figure 1C, Figure S2). The E/A identity of the nucleotide binding pair in GC1 is indicative of specificity towards GTP. Thus, we propose that GC1 and GC2 form a pseudo-heterodimer and function as a guanylate cyclase (Figure 1C). Similar to tmACs, one active and one degenerated active site is allocated at the dimer interface. However, the sequence of GC1 and GC2 is inverted in *Tg*ATPase_P_-GC, which means that, unlike tmAC, GC1 domain contributes to the nucleotide and transition state binding residues of the active site, whilst GC2 harbors two aspartates, crucial for metal ion binding.

### Overexpression and purification of recombinant GC1 and GC2 domains

With an objective to determine the functionality of *Tg*ATPase_P_-GC, we performed cloning of individual domains using the mRNA isolated from tachyzoites (RHΔ*ku80-hxgprt*) of *T. gondii*. We expressed the ORFs of GC1 (M^2850^-S^3244^), GC2 (M^3934^-Q^4242^) and GC1+GC2 (M^2850^-Q^4242^) in *E. coli* (Figure S3A). An overexpression of GC1 and GC2 as 6xHis-tagged in the M15 strain resulted in inclusion bodies, which we nevertheless purified by the virtue of a Ni-NTA column under denaturing conditions. Purified GC1 and GC2 exhibited an expected molecular weight of 44 and 34.2-kDa, respectively (Figure S3B). Our attempts to purify GC1+GC2 protein were futile however. To test the catalytic activity of purified GC1 and GC2 domains, we executed a guanylate cyclase assay. Both GC1 and GC2 were found inactive when tested separately or together. Further optimization of the assay yielded no detectable activity. We also examined whether GC1 and GC2 can function as adenylate cyclase using a bacterial complementation assay, as described by Karimova *et al*. (Karimova et al. 1998) (Figure S3C). GC1, GC2 and GC1+GC2 proteins were expressed in the BTH101 strain of *E. coli*, which is deficient in the adenylate cyclase activity and so unable to utilize maltose as a carbon source. The strain produced white colonies on MacConkey agar containing maltose, which would otherwise be red-colored upon induction of cAMP-dependent disaccharide catabolism. We observed that unlike the positive control (adenylate cyclase from *E. coli*), BTH101 strains expressing GC1, GC2 or GC1+GC2 produced only white colonies in each case (Figure S3C), which could either be attributed to suboptimal expression or a lack of adenylate cyclase activity in accord with the presence of signature residues defining the specificity for GTP in indicated domains (Figure S2).

### *Tg*ATPase_P_-GC is constitutively expressed in the plasma membrane at the apical pole

To gain insight into the endogenous expression and localization of *Tg*ATPase_P_-GC protein, we performed epitope tagging of the gene in tachyzoites of *T. gondii* (Figure 2A). The parental strain was transfected with a plasmid construct allowing 3’-insertional tagging of *Tg*ATPase_P_-GC with a hemagglutinin (HA) tag by single homologous crossover. The resulting transgenic strain (P_native_-*Tg*ATPase_P_-GC-HA_3’IT_-*Tg*Gra1_3’UTR_) encoded HA-tagged *Tg*ATPase_P_-GC under the control of its native promoter. Notably, the fusion protein localized predominantly at the apical end of the intracellularly proliferating parasites, as confirmed by its colocalization with *Tg*Gap45, a marker of the inner membrane complex (IMC) (Gaskins et al. 2004) (Figure 2A). The apical location of *Tg*ATPase_P_-GC-HA_3’IT_ was confirmed by its costaining with the IMC sub-compartment protein 1 (*Tg*ISP1) (Beck et al. 2010). Besides, we noted a significant expression of *Tg*ATPase_P_-GC-HA_3’IT_ outside the parasite periphery within the residual body, which has also been observed for several other proteins, such as Rhoptry Neck 4 (RON4) (Bradley et al. 2005). To assess the membrane location and predicted C-terminal topology of the protein, we stained extracellular parasites with α-HA antibody prior to and after detergent permeabilization of the parasite membranes (Figure 2B). The HA-staining was detected only after the permeabilization, indicating that C-terminus of *Tg*ATPase_P_-GC faces the parasite interior, as shown in the model (Figure 1B).

We then treated the extracellular parasites with α-toxin to split the plasma membrane (PM) from IMC to distinguish the distribution of *Tg*ATPase_P_-GC-HA_3’IT_ between both entities. By staining of tachyzoites with two markers, *i.e. Tg*Gap45 for the IMC and *Tg*Sag1 for the PM, we could show an association of *Tg*ATPase_P_-GC-HA_3’IT_ with the plasma membrane (Figure 2C). Making of a transgenic line encoding *Tg*ATPase_P_-GC-HA_3’IT_ also enabled us to evaluate its expression pattern by immunoblot analysis throughout the lytic cycle, which recapitulates the successive events of gliding motility, invasion of host cell, intracellular replication and egress leading to cell lysis. *Tg*ATPase_P_-GC is a bulky protein (477-kDa) with several transmembrane regions; hence it was not possible to resolve it by SDS-PAGE and transfer onto nitrocellulose membrane for staining. We therefore performed dot-blot analysis by loading protein samples directly onto an immunoblot membrane (Figure 2D-E). Unlike the parental strain (negative control), which showed only a faint (background) α-HA staining, we observed a strong signal in the *Tg*ATPase_P_-GC-HA_3’IT_-expressing strain (Figure 2D). Samples of transgenic strains collected at various periods embracing the entire lytic cycle of tachyzoites indicated a constitutive and stable expression of *Tg*ATPase_P_-GC-HA_3’IT_ (Figure 2E).

### *Tg*ATPase_P_-GC is essential for the parasite survival

Having established the expression profile and location, we next examined the physiological importance of *Tg*ATPase_P_-GC for tachyzoites of *T. gondii*. Our multiple attempts to delete the *Tg*ATPase_P_-GC gene by double homologous recombination were unrewarding suggesting its essentiality for the parasite (mortal phenotype). Hence, we utilized the aforementioned strain expressing *Tg*ATPase_P_-GC-HA_3’IT_ to trace the effect of gene disruption immediately after the plasmid transfection (Figure 3). To achieve this, we executed a CRISPR/Cas9-directed cleavage in *Tg*ATPase_P_-GC and then immunostained parasites at various periods to determine a time-elapsed loss of HA-signal (Figure 3A). Within a day of transfection, about 4% of vacuoles had lost the apical staining of *Tg*ATPase_P_-GC-HA_3’IT_ in contained parasites (Figure 3B). Parasites losing the HA-signal remained constant until the 1^st^ passage (24-40 h). However, their growth reduced gradually during the 2^nd^ passage (72-88 h) and fully seized by the 3^rd^ passage (120-136 h). The same assay also allowed us to quantify the replication rates of HA-negative parasites in relation to the HA-positive parasites by enumerating their numbers in intracellular vacuoles (Figure 3C). As expected, the fraction of small vacuoles comprising just 1 or 2 parasites was much higher in the nonexpression strain. Contrariwise, the progenitor strain encoding *Tg*ATPase_P_-GC-HA_3’IT_ showed predominantly a higher percentage of bigger vacuoles with 16-64 parasites. By 3^rd^ passage, we detected only single-parasite vacuoles in the mutant, demonstrating an essential role of *Tg*ATPase_P_-GC for the asexual reproduction.

**Figure 3.**
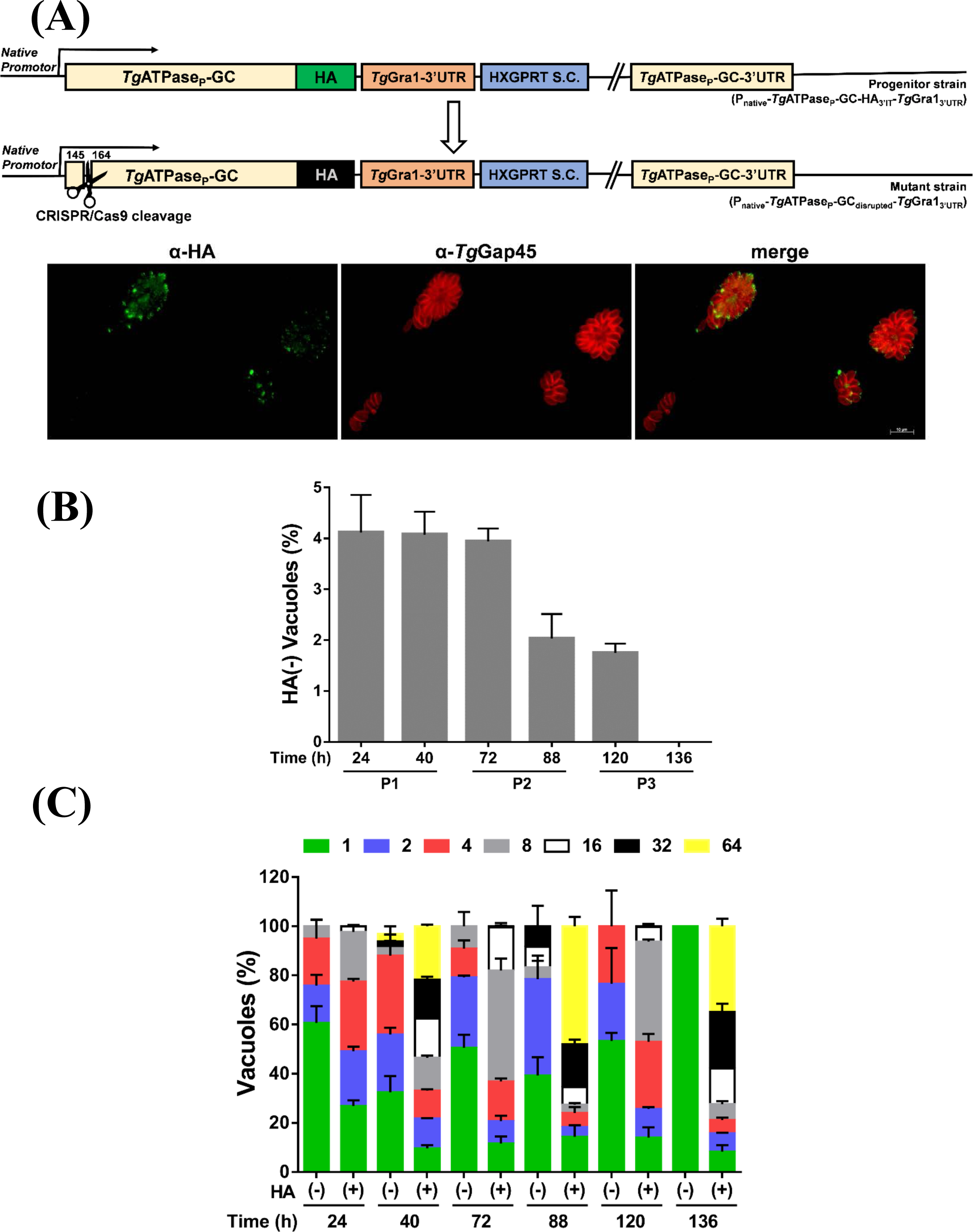
Genetic disruption of *Tg*ATPase_P_-GC is lethal for tachyzoites of *T. gondii*. **(A)** Scheme for the CRISPR/Cas9-mediated disruption of the gene in parasites expressing *Tg*ATPase_P_-GC-HA_3’IT_. The guide RNA was designed to target the nucleotides between 145 and 164 bp in the progenitor strain (P_native_-*Tg*ATPase_P_-GC-HA_3’IT_-*Tg*Gra1_3’UTR_). A representative image showing a loss of HA signal in some intracellular parasites replicating in the vacuole following a CRISPR/Cas9-cleavage in the indicated region of the *Tg*ATPase_P_-GC gene. Tachyzoites were transfected with the *pU6-TgATPase_P_-GC-sgRNA-Cas9* vector, and then stained with α-HA and α-*Tg*Gap45 antibodies at specified periods. **(B)** Quantitative graphical illustration of *Tg*ATPase_P_-GC-HA_3’IT_-disrupted mutant parasites from *panel a*. The HA-negative vacuoles harboring at least 2 parasites were scored at various time points during successive passages of culture (P1-P3). **(C)** The replication rates of the HA-positive and HA-negative tachyzoites including single parasites, as evaluated by immunostaining (*panel a*). Approximately, 500-600 vacuoles from three independent assays were scored with S.E.M by enumerating the number of parasites per vacuole.

### Genetic knockdown of cGMP synthesis impairs the lytic cycle

Although an indispensable nature of *Tg*ATPase_P_-GC for tachyzoites could be recognized, the above strategy did not yield us a clonal mutant line for in-depth biochemical and phenotypic analyses due to an eventually lethal phenotype. To circumvent this issue, we first endeavored tagging the gene with an auxin-inducible degradation domain to make a conditional mutant, as described elsewhere (Long et al. 2017). The strategy did not result into viable drug-resistant parasites following transgenic selection. We therefore engineered yet other parasite strain expressing *Tg*ATPase_P_-GC-HA_3’IT_, in which the native 3’UTR of the gene was flanked with the Cre/loxP sites (Figure 4A). Cre recombinase-mediated excision of the 3’UTR combined with a negative selection, as reported earlier (Brecht et al. 1999), permitted a destabilization of *Tg*ATPase_P_-GC. Genomic screening using the specific primers confirmed a successful generation of the clonal mutant lines (P_native_-*Tg*ATPase_P_-GC-HA_3’IT_-3’UTR_excised_), which yielded a 2.2-kb amplicon as opposed to 5.2-kb in the progenitor (P_native_-*Tg*ATPase_P_-GC-HA_3’IT_-3’UTR_floxed_) (Figure 4B). Immunoblots of selected clones showed an evident repression of the protein (Figure 4C) that was ascertained by a loss of HA-staining in immunofluorescence assays (Figure 4D).

**Figure 4.**
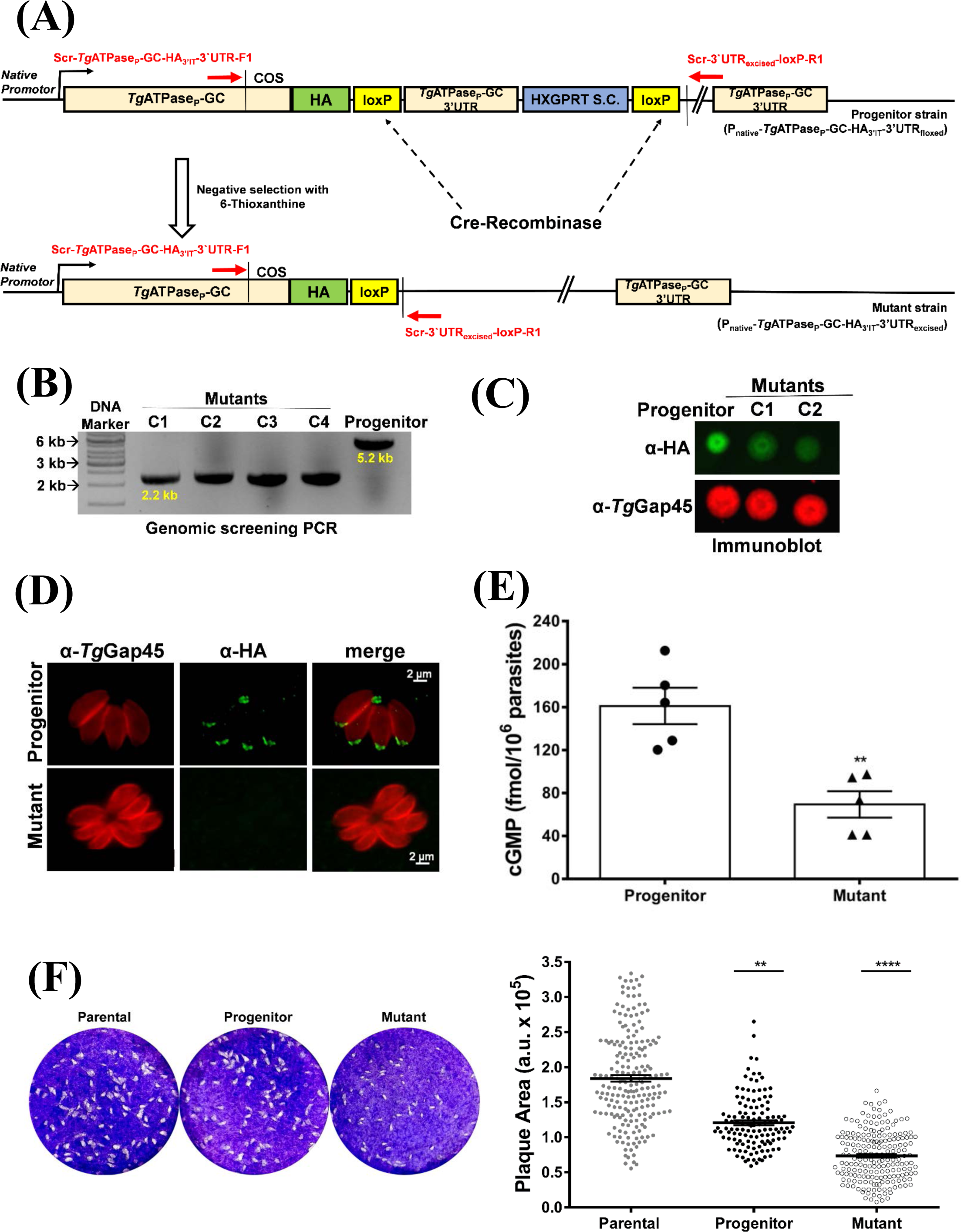
Cre recombinase-mediated downregulation of cGMP synthesis impairs the lytic cycle of *T. gondii*. **(A)** Schematics for making the parasite mutant (P_native_-*Tg*ATPase_P_-GC-HA_3’IT_-3’UTR_excised_). A vector expressing Cre recombinase was transfected into the progenitor strain (P_native_-*Tg*ATPase_P_-GC-HA_3’IT_-3’UTR_floxed_), in which 3’UTR of the *Tg*ATPase_P_-GC gene was flanked with two Cre/loxP sites. Parasites transfected with a Cre-recombinase expressing vector were negatively selected by 6-thioxanthine for the loss of HXGPRT selection cassette (S.C.). **(B)** Genomic screening of the *Tg*ATPase_P_-GC mutant confirming Cre-mediated excision of 3’UTR and HXGPRT. Primers indicated as red arrows in *panel a* were used to screen the gDNAs isolated from four different mutant clones (C1-C4) along with the progenitor strain. **(C)** Immunoblot depicting a repression of *Tg*ATPase_P_-GC-HA_3’IT_ in parasites with excised 3’UTR in comparison to the progenitor strain. Tachyzoites (10^7^) were subjected to dot blot analysis using α-HA, and α-*Tg*Gap45 (loading control) antibodies. **(D)** Immunostaining of the mutant (*Tg*ATPase_P_-GC-HA_3’IT_-3’UTR_excised_) and progenitor parasites revealing a loss of HA-signal in the former strain. Parasites were stained with α-HA and α-*Tg*Gap45 antibodies 24 h post infection. **(E)** Changes in total cGMP level of the mutant compared to the progenitor strain. Fresh syringe-released tachyzoites (5×10^6^) were subjected to ELISA-based cGMP measurements. **(F)** Plaque assays of the *Tg*ATPase_P_-GC mutant along with its progenitor and parental strains. The dotted white areas and blue staining signify the parasite plaques and host monolayers, respectively (*left*). The area of each plaque, calculated in relative arbitrary units (a. u.) using ImageJ program, reveals the overall growth fitness of strains. 150-200 plaques of each strain were evaluated *(right)*. The *panel e* and *panel f* show data from 5 and 3 independent assays, respectively with S.E.M. (**, p ≤0.01; ****, p ≤0.0001).

Next, we evaluated if a knockdown of *Tg*ATPase_P_-GC translated into a declined synthesis of cGMP by the parasite. Indeed, we measured about 50% decline in the steady-state levels of cGMP in the mutant corresponding to decay in the protein level (Figure 4E). We then measured the comparative fitness of the mutant, progenitor and parental (RHΔ*ku80-hxgprt^−^*) strains by plaque assays (Figure 4F). As anticipated, the mutant exhibited a significant ~55% reduction in plaque area when compared to the parental, which correlated with its residual expression of *Tg*ATPase_P_-GC as well as with its cGMP levels (Figure 4C-E). Interestingly, the progenitor strain itself showed an about 30% impairment that is likely due to epitope-tagging and introduction of Cre/loxP site between the last exon of the gene and its 3’UTR (Figure 4A). Together with above results, these data show that *Tg*ATPase_P_-GC functions as a guanylate cyclase, and its catalytic activity is indispensable for the lytic cycle.

### *Tg*ATPase_P_-GC regulates multiple events during the lytic cycle

The availability of a functional mutant prompted us to study the importance of *Tg*ATPase_P_-GC for discrete steps of lytic cycle including host-cell invasion, intracellular proliferation, egress and gliding motility (Figure 5). The replication assay at two different time points (24 h and 40 h) revealed a significantly higher fraction of smaller vacuoles in the mutant, particularly in early cultures (24 h) (Figure 5A). The effect was alleviated and somewhat negligible in late-stage cultures (40 h). We quantified ~30% decline in invasion efficiency of the mutant (down from 80% to 53%) compared to the parental (RHΔ*ku80-hxgprt^−^*) strain (Figure 5B). The effect of enzyme downregulation was more pronounced in egress assay, in which the mutant showed >50% decline in natural egress of mature parasites at 40 h and 48 h post-infection when compared to the parental strain (Figure 5C). The egress defect was not apparent however upon prolonged incubation (64 h). Since invasion and egress are mediated by gliding motility (Frénal et al. 2017), we tested our mutant for the latter phenotype. Indeed, we determined that the average motile fraction of the mutant was reduced by more than half, and the trail lengths of moving tachyzoites were remarkably shorter (~18 µm) than the progenitor strain (~50 µm) (Figure 5D). Not least, as seen in the plaque assay (Figure 4F), we found a steady albeit not significant decline in the replication, invasion and egress of the progenitor in relation to the parental strain, further confirming a correlation across all phenotypic assays.

**Figure 5.**
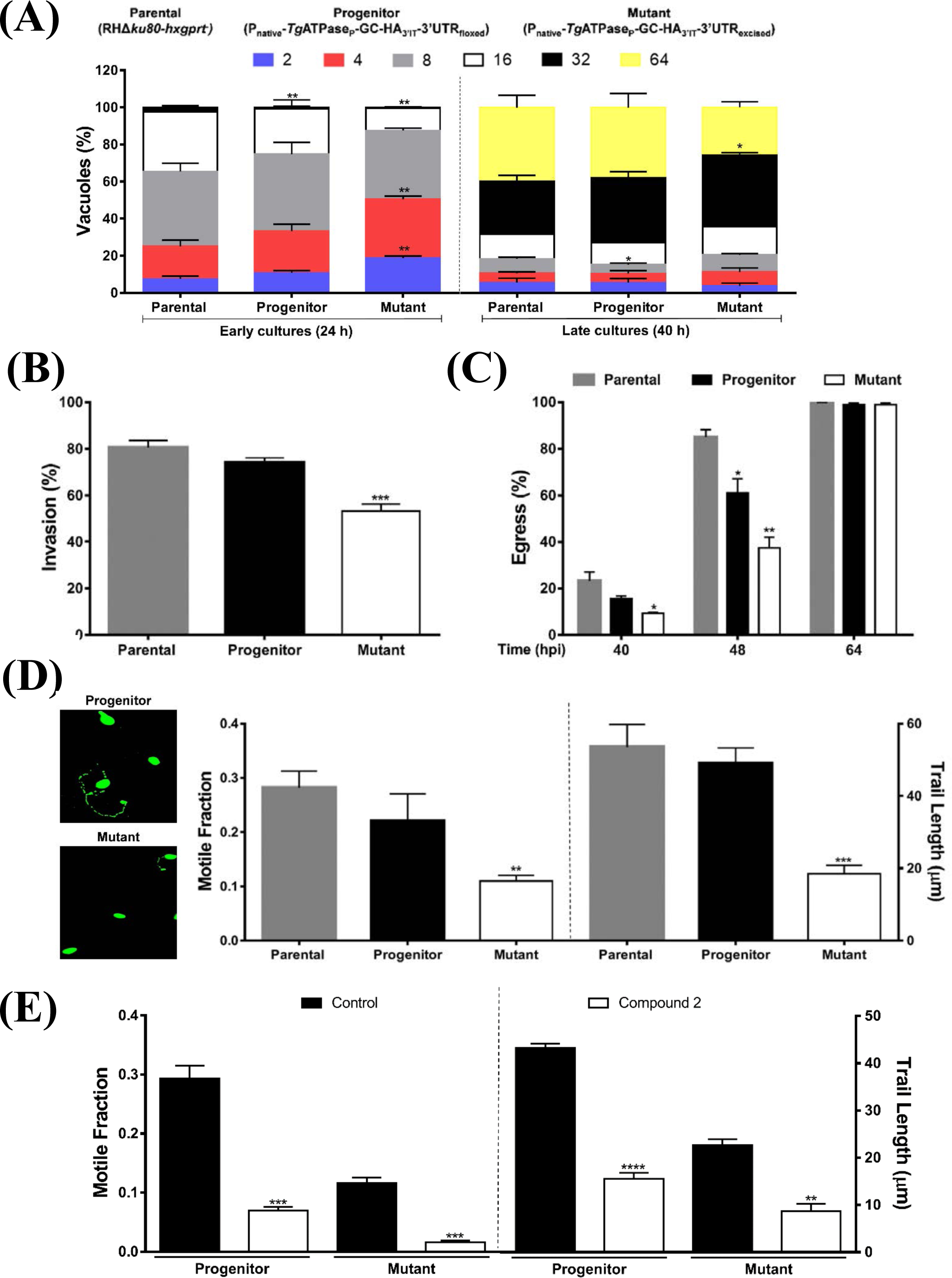
Cyclic GMP signaling governs the key events during the lytic cycle of *T. gondii*. **(A-D)** In-depth phenotyping of the *Tg*ATPase_P_-GC mutant (P_native_-*Tg*ATPase_P_-GC-HA_3’IT_-3’UTR_excised_), its progenitor (P_native_-*Tg*ATPase_P_-GC-HA_3’IT_-3’UTR_floxed_) and parental (RHΔ*ku80-hxgprt^−^*) strains. The intracellular replication **(A)**, host-cell invasion **(B)**, parasite egress **(C)** and gliding motility **(D)** were assessed using standard phenotyping methods, as described in *methods* section. The progenitor and mutant strains were generated as shown in Figure 2A and Figure 4A, respectively. The replication rates were analyzed at two different time points (24 h and 40 h) by scoring the parasite numbers in a total of 500-600 vacuoles following their staining with α-*Tg*Gap45 antibody (n= 4 assays). Invasion and egress rates were calculated by dual-staining with α-*Tg*Gap45 and α-*Tg*Sag1 antibodies as described in *methods* section. Approximately, 1000 parasites for each strain from 4 assays were examined to estimate the invasion efficiency (1 h incubation). The natural egress of tachyzoites was measured after different periods (40 h, 48 h and 64 h) by scoring 500-600 vacuoles for each strain from 3 assays. To estimate the motility, immunofluorescence images stained with α-*Tg*Sag1 antibody were analyzed for the motile fraction (about 500 parasites of each strain), and 100-120 trail lengths per strain were measured from 3 assays. **(E)** The effect of PKG inhibitor compound 2 (2 µM) on the motility of *Tg*ATPase_P_-GC mutant and its progenitor strain (n= 3 assays). A total of 100 trails in the progenitor and only 15 trails of the mutant (due to severe defect) were measured. Numerical values in graphs from *panel a-e* show the means with S.E.M. Statistical significance in individual assays were tested by comparing the progenitor and mutant strains with the parental strain (*, p ≤0.05; **, p ≤0.01; ***, p ≤0.001; ****, p ≤0.0001).

A partial phenotype prompted us to inhibit the residual cGMP signaling pharmacologically *via* PKG in the *Tg*ATPase_P_-GC mutant. We utilized compound 2 (C2), which has been shown to block mainly *Tg*PKG, but also calcium-dependent protein kinase 1 (Donald et al. 2006). As rationalized, C2 treatment subdued the gliding motility of the mutant as well as of the progenitor strain (Figure 5E). The impact of C2 was accentuated in both strains, which is likely a cumulative effect of genetic repression and drug inhibition. Inhibition was stronger in the *Tg*ATPase_P_-GC mutant than the progenitor strain, which can be attributed to inhibition of the residual cGMP signaling in the knockdown strain.

### Phosphodiesterase inhibitors can rescue defective phenotypes of *Tg*ATPase_P_-GC mutant

To further validate our findings, we deployed two inhibitors of cGMP-specific PDEs, zaprinast and BIPPO, which are known to inhibit parasite enzymes along with human PDE5 and PDE9, respectively (Howard et al. 2015, Yuasa et al. 2005). We reasoned that pharmacological elevation of cGMP should assuage the phenotypic defects caused by deficiency of guanylate cyclase in the mutant. As shown (Figure S4A), both drugs led to a dramatic increase in the motile fraction and trail lengths of the progenitor and *Tg*ATPase_P_-GC mutant. The latter parasites were as competent as the former after the drug exposure. A similar restoration of phenotype in the mutant was detected in egress assays; the effect of BIPPO was much more pronounced than zaprinast however (Figure S4B). BIPPO was almost 2x more efficient than zaprinast leading to egress of almost all parasites. In contrast to motility and egress, a treatment of BIPPO and zaprinast resulted in a surprisingly divergent effect on the invasion rates of the two strains (Figure S4C). BIPPO exerted an opposite effect, *i.e*. a reduction in invasion of the progenitor and mutant. Impairment was stronger in the earlier strain; hence we noticed a complete reversal of the phenotype when compared to control samples. A fairly similar effect was seen with zaprinast; though, it was much less potent than BIPPO, as implied previously (Howard et al 2015). These observations can be attributed to differential elevation of cGMP (above certain threshold) caused by PDE-inhibitors, which inhibits the host-cell invasion, but promotes the parasite motility and egress.

### Genetic knockdown of *Tg*PKG phenocopies the attenuation of *Tg*ATPase_P_-GC

To consolidate aforesaid work, we implemented the same genomic-tagging, knockdown and phenotyping approach to *Tg*PKG, which is the major mediator of cGMP signaling initiated by *Tg*ATPase_P_-GC (Figure S5, Figure 6). Briefly, a transgenic strain expressing *Tg*PKG with a C-terminal HA-tag under the control of endogenous regulatory elements and floxed 3’UTR was generated by 3’-insertional tagging strategy (Figure S5A). The resultant progenitor strain (P_native_-*Tg*PKG-HA_3’IT_-3’UTR_floxed_), subjected to immunostaining, revealed an expression of *Tg*PKG-HA_3’IT_ in plasma membrane and cytoplasm of intracellular parasites, verifying earlier reports (Brown et al. 2017, Donald and Liberator 2002, Gurnett et al. 2002) (Figure 6A). We then executed Cre-mediated knockdown of *Tg*PKG-HA_3’IT_ by excising the loxP-flanked 3’UTR in the progenitor strain (Figure S5A). As intended, genomic screening with specified primers yielded 1.9-kb amplicons in the isolated clonal mutants (P_native_-*Tg*PKG-HA_3’IT_-3’UTR_excised_) as opposed to a 4.8-kb band in the progenitor strain (P_native_-*Tg*PKG-HA_3’IT_-3’UTR_floxed_), which confirmed the excision of 3’UTR (Figure S5B). A repression of *Tg*PKG-HA_3’IT_ protein was validated by immunofluorescence (Figure 6A) and immunoblot assays (Figure S5C). As shown before (Brown, Long and Sibley 2017, Donald and Liberator 2002, Gurnett, Liberator, Dulski, Salowe, Donald, Anderson, Wiltsie, Diaz, Harris and Chang 2002), we also observed two isoforms of *Tg*PKG (encoded by a single gene) in the progenitor strain. Excision of 3’UTR resulted in a knockdown of both isoforms (112-kDa and 135-kDa) in the clonal mutant strains of *Tg*PKG-HA_3’IT_ (Figure S5C).

**Figure 6.**
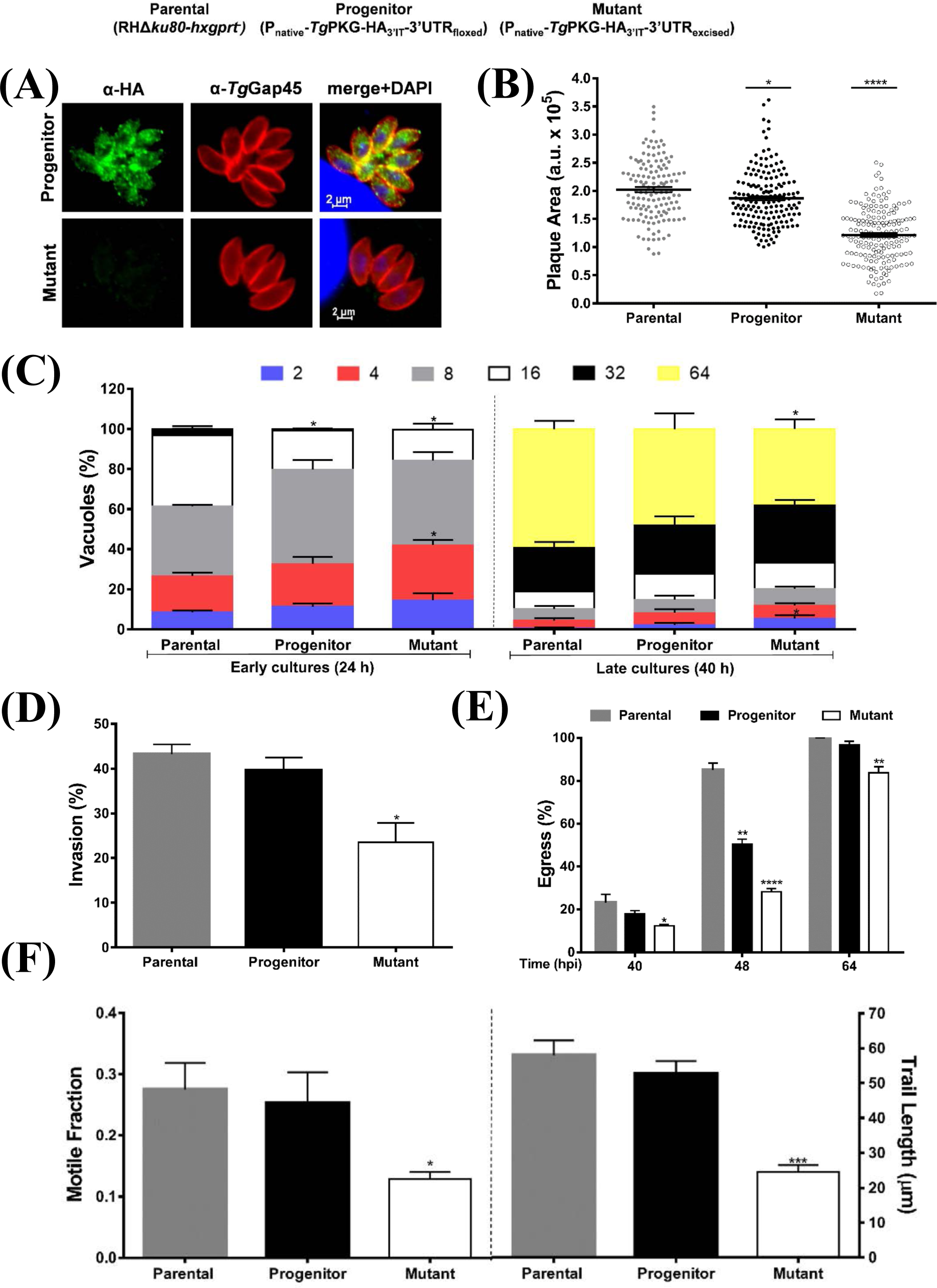
Mutagenesis of *Tg*PKG recapitulates the phenotype of the *Tg*ATPase_P_-GC mutant. **(A-F)** Phenotyping of the *Tg*PKG mutant (P_native_-*Tg*PKG-HA_3’IT_-3’UTR_excised_) in comparison to its progenitor (P_native_-*Tg*PKG-HA_3’IT_-3’UTR_floxed_) and parental (RHΔ*ku80-hxgprt^−^*) strains. For making of the mutant, refer to Figure S5. **(A)** Immunofluorescence images demonstrating the subcellular localization of *Tg*PKG-HA_3’IT_ in cytomembranes of the progenitor strain, and its downregulation in the mutant. Intracellular parasites (24 h post-infection) were stained with α-HA and α-*Tg*Gap45 antibodies, as described in *methods*. The merged image shows DAPI-stained host and parasite nuclei in blue. **(B)** Plaque assay revealing comparative growth of the mutant, progenitor and parental strains. The plaque area is shown as arbitrary units (a. u.), was measured by ImageJ software. A total of 140-170 plaques for each strain were scored from 3 assays. **(C)** Replication rates of indicated strains during early (24 h) and late (40 h) cultures. Intracellular tachyzoites proliferating in their vacuoles were immunostained with α-*Tg*Gap45 antibody. The number of tachyzoites per vacuole were counted from 400-500 vacuoles for each strain (n= 3 assays). **(D-E)** Invasion and egress of the *Tg*PKG mutant along with the parental and progenitor parasites, as judged by dual-color staining. Intracellular tachyzoites were immunostained red using α-*Tg*Gap45 antibody, while extracellular ones appeared two-colored (red and green) stained with both α-*Tg*Gap45 and α-*Tg*Sag1 antibodies. In total, 1000-1200 parasites were evaluated to score the invasion rate of each strain (n= 5 assays). The percentage of ruptured vacuoles at indicated periods was determined by observing 400-500 vacuoles for each strain from 3 experiments. **(F)** The motile fraction and trail lengths of the indicated parasite strains. About 600 parasites were analyzed for the motile fraction, and 100 trail lengths were measured using ImageJ software, following immunostaining with α-*Tg*Sag1 (n= 3 assays). Numerical values in all graphs from the *panel b-f* show the means with S.E.M. Statistical significance in individual assays was measured by comparing the progenitor and mutant strains against the parental strain (*, p ≤0.05; **, p ≤0.01; ***, p ≤0.001; ****, p ≤0.0001).

Cre recombinase-mediated destabilization of *Tg*PKG protein led to an analogous inhibition of the parasite growth in plaque assays (Figure 6B). The plaques were much smaller in the mutant; yet again, analogous to the *Tg*ATPase_P_-GC mutant (Figure 4F), replication of the *Tg*PKG mutant was moderately down, as judged by a smaller fraction of bigger vacuoles containing 32 or 64 parasites in early (24 h) and late (40 h) cultures (Figure 6C). We also scored a noteworthy invasion defect in the *Tg*PKG mutant (Figure 6D). In accord, the mutant exhibited a defective egress at all tested time points (Figure 6E). The motile fraction dropped by almost 50% in the mutant, and trail lengths were accordingly shorter (~24 µm) compared to the control strains (~55 µm) (Figure 6F). Not least, a treatment of compound 2 yielded further declines in the motile fraction and trail lengths of the mutant and its progenitor strain (Figure S6). The impact of C2 was somewhat stronger in the mutant, and none of the two strains exhibited a complete inhibition, once more resonating with our data on the *Tg*ATPase_P_-GC mutant (Figure 5E). These data clearly show that repression of *Tg*ATPase_P_-GC or *Tg*PKG protein imposes nearly identical phenotypic defects on the lytic cycle of *T. gondii*.

## DISCUSSION

In this study, we revealed that *T. gondii* encodes an unusual evidently multifunctional protein *Tg*ATPase_P_-GC, which is primarily localized in the plasma membrane at the apical pole of the tachyzoite stage, while *Tg*PKG is located in cytomembranes (Brown, Long and Sibley 2017)(this study). Using the same approach, we found that *Tg*ATPase_P_-GC and *Tg*PKG are expressed through the lytic cycle, suggesting a post-translational control of signaling. Both proteins are critical for a successful progression of the lytic cycle, where they control motility-dependent egress and invasion of the parasite. Essentiality of PKG for the asexual lifecycle in coccidian parasites (*T. gondii* and *E. tenella*) was first revealed by a chemical-genetic approach (Donald et al. 2002). Successive works on *T. gondii* have endorsed a vital requirement of *Tg*PKG for its asexual reproduction by various methods (Brown, Long and Sibley 2017, Donald and Liberator 2002, Lourido et al. 2012, Sidik et al. 2014). Lately, it was shown that of the two alternatively-translated isoforms the one located in the plasma membrane is essential to govern the PKG-dependent events, whereas the other residing in the cytoplasm is dispensable (Brown, Long and Sibley 2017). Likewise, PKG also controls the hepatic and erythrocytic development of *Plasmodium* species (Falae et al. 2010, Taylor et al. 2010). Characterization of *Tg*ATPase_P_-GC herein, as the enzyme involved in cGMP synthesis, imparts a central piece of signaling conundrum in *T. gondii*. Its noticeably predicted multi-functionality breeds several premises, advocating further explorations to dissect the functional repurposing and evolution of cGMP cascade in protozoans.

The membrane topology, subcellular distribution, structural modeling and predictive functions of *Tg*ATPase_P_-GC are strikingly distinct from the particulate GCs (pGCs) of mammalian cells, especially due to the possession of manifold ATPase-like motifs in the N-terminal extension. The malaria parasite *Plasmodium* also contains two guanylate cyclases (GCα, GCβ), both of which are marked by the presence of a P-type ATPase-like domain (Baker 2004, Hopp, Bowyer and Baker 2012). Of note is also the fact that this topology is shared by members of another alveolate phylum, namely Ciliophora (*e.g. Paramecium*, *Tetrahymena*) (Linder et al. 1999), which exhibit an entirely different lifestyle. Sequence analysis indicated that the guanylate cyclase domain of this seemingly multifaceted protein is not related to the mammalian guanylate cyclases; instead they seem to have evolved from the class III adenylate cyclases. A guanylate cyclase having two cyclase domains with a similar topology also occurs in a member of Amoebozoa (*Dictyostelium*), which is devoid of the ATPase-like domain though (Roelofs et al. 2001). GC1 and GC2 have probably evolved by gene duplication event causing degeneration of the unused second regulatory binding site, as suggested for tmACs (Linder 2005, Steegborn 2014, Tesmer et al. 1997). Consequently, *Tg*ATPase_P_-GC forms only one pseudo-symmetric catalytic center by dimerization of GC1 and GC2 domains, in contrast to homodimer formation as in mammalian pGCs (Linder 2005, Linder and Schultz 2003, Steegborn 2014). Other distinguished features of pGCs, the extracellular ligand binding and regulatory kinase-homology domains, are also not found in the protozoan counterparts, adding to evolutionary specialization of cGMP signaling.

Unlike the C-terminal guanylate cyclase, the function of N-terminal ATPase motifs remains enigmatic; it resembles the type-IV P-type ATPases in most alveolate GCs (Baker et al. 2017, Carucci et al. 2000, Kenthirapalan et al. 2016, Linder et al. 1999). As shown, *Tg*ATPase_P_-GC harbors at least four motifs, possibly serving distinct purposes in *T. gondii*. Given the type of residues in *Tg*ATPase_P_-GC and existing literature on the regulation of lytic cycle, we postulate phospholipid flipping and cation transport as the two key functions of the N-terminal domains. Phosphatidic acid and cation signaling/homeostasis, for example, have been shown to influence the events of gliding motility and associated protein secretion, which in turn drive the egress and invasion events (Bullen et al. 2016, Endo and Yagita 1990, Frénal et al. 2017, Rohloff et al. 2011). There is no evidence how lipid and cation-dependent pathways embrace each other. It is therefore tempting to speculate an integrative nodal role of *Tg*ATPase_P_-GC in asymmetric distribution of phosphatidic acid in the membrane leaflets and cation flux (*e.g*. Ca^2+^, K^+^, Na^+^) across the plasma membrane leaflets. At this point, intramolecular coordination of ATPase and guanylate cyclase in *Tg*ATPase_P_-GC also remains unclear. Whether the GC domains can influence P-type ATPase motifs, or *vice versa*, seem equally plausible. A mechanistic understanding of this mega-protein shall reveal the master regulation of the lytic cycle.

Similar to *Tg*ATPase_P_-GC, PKGs in protists have also diverged from mammalian equivalents. Although, protozoan PKGs have kept the fundamental features of the protein kinase family, they differ from other known PKG sequences in carrying more than two cGMP-binding sites. Analysis of *Toxoplasma*, *Eimeria* and *Plasmodium* PKGs revealed that apicomplexan parasites contain three high probability (stated as A-B-D to show the location on the gene) and an additional low probability (C) cyclic nucleotide binding site, while *Paramecium* PKG harbors two high (A and B) and one low probability (C) site (Baker and Deng 2005). The phosphorylation of a threonine residue in the catalytic domain of mammalian PKG was reported as essential for the enzyme activity (Feil et al. 1995); this site is either auto-phosphorylated or targeted by other kinases, which seems not be the case in protozoan PKG, as demonstrated for *Pf*PKG (Baker and Deng 2005, Feil, Kellermann and Hofmann 1995). It is also worth mentioning that PKG-independent effectors of cGMP, *i.e*. nucleotide-gated ion channels as reported in mammalian cells (Lucas et al. 2000, MacFarland 1995, Pilz and Casteel 2003), could not be identified in the genomes of alveolates, suggesting a rather linear transduction of cGMP pathways in the latter organisms. Besides, a conserved design of cGMP signaling in alveolates having otherwise diverse lifestyles entails a convoluted repurposing of cGMP signaling within the kingdom. Last but not least, a divergent origin and essential requirement of the cGMP cascade in the parasitic protists can be exploited to selectively inhibit their reproduction.

## Supporting information

## Abbreviations

cGMP: cyclic guanosine monophosphate
GC: guanylate cyclase
PKG: cGMP-dependent protein kinase
PDE: phosphodiesterase
HFF: human foreskin fibroblast
MOI: multiplicity of infection
COS: crossover sequence
IT: insertional tagging
HXGPRT: hypoxanthine-xanthine-guanine-phosphoribosyltransferase
GOI: gene of interest
UTR: untranslated region
PFA: paraformaldehyde
IPTG: Isopropyl β-D-1-thiogalactopyranoside

## Acknowledgment

We thank Grit Meusel (Humboldt University) for her technical assistance. In addition, we are grateful to the parasitology community for sharing critical reagents. This work was supported by research grants (GU1100/7-1, GRK2046) and Heisenberg program fellowship (GU1100/8-1), awarded to NG by German Research Foundation (DFG). Financial support to ÖGE was provided through Elsa-Neumann-Scholarship by the state of Berlin.

## Author contributions statement

ÖGE performed the experiments; US contributed new sequence analysis and modeling skills; NG coordinated the project; ÖGE and NG, wrote the manuscript. All coauthors reviewed and approved the manuscript.

## REFERENCES

Arroyo-Olarte RD, Brouwers JF, Kuchipudi A, Helms JB, Biswas A, Dunay IR, Lucius R, and Gupta N. (2015). Phosphatidylthreonine and lipid-mediated control of parasite virulence. PLoS Biology.13:e1002288.

Baker D. (2004). Adenylyl and guanylyl cyclases from the malaria parasite Plasmodium falciparum. IUBMB life.56:535–540.

Baker DA, Deng W. (2005). Cyclic GMP-dependent protein kinases in protozoa. Front Biosci.10:1229–1238.

Baker DA, Drought LG, Flueck C, Nofal SD, Patel A, Penzo M, and Walker EM. (2017). Cyclic nucleotide signalling in malaria parasites. Open Biology.7:170213.

Beavo JA. (1995). Cyclic nucleotide phosphodiesterases: functional implications of multiple isoforms. Physiological Reviews.75:725–748.

Beck JR, Rodriguez-Fernandez IA, De Leon JC, Huynh M-H, Carruthers VB, Morrissette NS, and Bradley PJ. (2010). A novel family of *Toxoplasma* IMC proteins displays a hierarchical organization and functions in coordinating parasite division. PLoS Pathogens.6:e1001094.

Biftu T, Feng D, Ponpipom M, Girotra N, Liang G-B, Qian X, Bugianesi R, Simeone J, Chang L, and Gurnett A. (2005). Synthesis and SAR of 2, 3-diarylpyrrole inhibitors of parasite cGMP-dependent protein kinase as novel anticoccidial agents. Bioorganic & Medicinal Chemistry Letters.15:3296–3301.

Blume M, Rodriguez-Contreras D, Landfear S, Fleige T, Soldati-Favre D, Lucius R, and Gupta N. (2009). Host-derived glucose and its transporter in the obligate intracellular pathogen *Toxoplasma gondii* are dispensable by glutaminolysis. Proceedings of the National Academy of Sciences.106:12998–13003.

Bradley PJ, Ward C, Cheng SJ, Alexander DL, Coller S, Coombs GH, Dunn JD, Ferguson DJ, Sanderson SJ, and Wastling JM. (2005). Proteomic analysis of rhoptry organelles reveals many novel constituents for host-parasite interactions in Toxoplasma gondii. Journal of Biological Chemistry. 280:34245–34258.

Brecht S, Erdhart H, Soete M, and Soldati D. (1999). Genome engineering of *Toxoplasma gondii* using the site-specific recombinase Cre. Gene.234:239–247.

Brown KM, Long S, and Sibley LD. (2017). Plasma Membrane Association by N-Acylation governs PKG function in *Toxoplasma gondii*. mBio.8:e00375–00317.

Bublitz M, Morth JP, and Nissen P. (2011). P-type ATPases at a glance. Journal of Cell Science.124:2515–2519.

Bullen HE, Jia Y, Yamaryo-Botté Y, Bisio H, Zhang O, Jemelin NK, Marq J-B, Carruthers V, Botté CY, and Soldati-Favre D. (2016). Phosphatidic acid-mediated signaling regulates microneme secretion in *Toxoplasma*. Cell Host & Microbe.19:349–360.

Carucci DJ, Witney AA, Muhia DK, Warhurst DC, Schaap P, Meima M, Li J-L, Taylor MC, Kelly JM, and Baker DA. (2000). Guanylyl cyclase activity associated with putative bifunctional integral membrane proteins in *Plasmodium falciparum*. Journal of Biological Chemistry.275:22147–22156.

Donald RG, Allocco J, Singh SB, Nare B, Salowe SP, Wiltsie J, and Liberator PA. (2002). *Toxoplasma gondii* cyclic GMP-dependent kinase:Chemotherapeutic targeting of an essential parasite protein kinase. Eukaryotic Cell.1:317–328.

Donald RG, Carter D, Ullman B, and Roos DS. (1996). Insertional tagging, cloning, and expression of the *Toxoplasma gondii* hypoxanthine-xanthine-guanine phosphoribosyltransferase gene. Use as a selectable marker for stable transformation. The Journal of Biological Chemistry. Jun 14;271:14010–14019. Epub 1996/06/14.

Donald RG, and Liberator PA. (2002). Molecular characterization of a coccidian parasite cGMP dependent protein kinase. Molecular and Biochemical Parasitology.120:165–175.

Donald RG, Zhong T, Wiersma H, Nare B, Yao D, Lee A, Allocco J, and Liberator PA. (2006). Anticoccidial kinase inhibitors: Identification of protein kinase targets secondary to cGMP-dependent protein kinase. Molecular and Biochemical Parasitology.149:86–98.

Dubremetz JF, Achbarou A, Bermudes D, and Joiner KA. (1993). Kinetics and pattern of organelle exocytosis during *Toxoplasma gondii*/host-cell interaction. Parasitology Research.79:402–408.

Endo T, and Yagita K. (1990). Effect of extracellular ions on motility and cell entry in *Toxoplasma gondii*. Journal of Eukaryotic Microbiology.37:133–138.

Falae A, Combe A, Amaladoss A, Carvalho T, Menard R, and Bhanot P. (2010). Role of *Plasmodium berghei* cGMP-dependent protein kinase in late liver stage development. Journal of Biological Chemistry.285:3282–3288.

Feil R, Kellermann J, and Hofmann F. (1995). Functional cGMP-dependent protein kinase is phosphorylated in its catalytic domain at threonine-516. Biochemistry.34:13152–13158.

Fox BA, Ristuccia JG, Gigley JP, and Bzik DJ. (2009). Efficient gene replacements in *Toxoplasma gondii* strains deficient for nonhomologous end joining. Eukaryot Cell. Apr;8:520–529. Epub 2009/02/17.

Frénal K, Dubremetz J-F, Lebrun M, and Soldati-Favre D. (2017). Gliding motility powers invasion and egress in Apicomplexa. Nature Reviews Microbiology.15:645.

Gajria B, Bahl A, Brestelli J, Dommer J, Fischer S, Gao X, Heiges M, Iodice J, Kissinger JC, Mackey AJ, et al. (2008). ToxoDB: an integrated *Toxoplasma gondii* database resource. Nucleic Acids Research. Jan;36:D553–556. Epub 2007/11/16.

Gaskins E, Gilk S, DeVore N, Mann T, Ward G, and Beckers C. (2004). Identification of the membrane receptor of a class XIV myosin in *Toxoplasma gondii*. The Journal of Cell Biology.165:383–393.

Gould MK, and de Koning HP. (2011). Cyclic-nucleotide signalling in protozoa. FEMS Microbiology Reviews.35:515–541.

Govindasamy K, Jebiwott S, Jaijyan D, Davidow A, Ojo K, Van Voorhis W, Brochet M, Billker O, and Bhanot P. (2016). Invasion of hepatocytes by *Plasmodium* sporozoites requires cGMP□dependent protein kinase and calcium dependent protein kinase 4. Molecular Microbiology.102:349–363.

Gupta N, Zahn MM, Coppens I, Joiner KA, and Voelker DR. (2005). Selective disruption of phosphatidylcholine metabolism of the intracellular parasite *Toxoplasma gondii* arrests its growth. Journal of Biological Chemistry.280:16345–16353.

Gurnett AM, Liberator PA, Dulski PM, Salowe SP, Donald RG, Anderson JW, Wiltsie J, Diaz CA, Harris G, and Chang B. (2002). Purification and molecular characterization of cGMP-dependent protein kinase from apicomplexan parasites a novel chemotherapeutic target. Journal of Biological Chemistry.277:15913–15922.

Hall CL, and Lee VT. (2018). Cyclic□di□GMP regulation of virulence in bacterial pathogens. Wiley Interdisciplinary Reviews: RNA. 9.

Hofmann K, and Stoffel W. (1993). TMbase - A database of membrane spanning proteins segments. Biol. Chem. Hoppe-Seyler.374:166.

Hopp CS, Bowyer PW, and Baker DA. (2012). The role of cGMP signalling in regulating life cycle progression of *Plasmodium*. Microbes and Infection.14:831–837.

Howard BL, Harvey KL, Stewart RJ, Azevedo MF, Crabb BS, Jennings IG, Sanders PR, Manallack DT, Thompson PE, and Tonkin CJ. (2015). Identification of potent phosphodiesterase inhibitors that demonstrate cyclic nucleotide-dependent functions in apicomplexan parasites. ACS Chemical Biology.10:1145–1154.

Huynh MH, and Carruthers VB. (2009). Tagging of endogenous genes in a *Toxoplasma gondii* strain lacking Ku80. Eukaryot Cell. Apr;8:530–539. Epub 2009/02/17.

Käll L, Krogh A, and Sonnhammer EL. (2004). A combined transmembrane topology and signal peptide prediction method. Journal of Molecular Biology.338:1027–1036.

Karimova G, Pidoux J, Ullmann A, and Ladant D. (1998). A bacterial two-hybrid system based on a reconstituted signal transduction pathway. Proceedings of the National Academy of Sciences.95:5752–5756.

Kenthirapalan S, Waters AP, Matuschewski K, and Kooij TW. (2016). Functional profiles of orphan membrane transporters in the life cycle of the malaria parasite. Nature communications.7:10519.

Kong P, Lehmann MJ, Helms JB, Brouwers JF, and Gupta N. (2018). Lipid analysis of *Eimeria* sporozoites reveals exclusive phospholipids, a phylogenetic mosaic of endogenous synthesis, and a host-independent lifestyle. Cell Discovery.4:24.

Letunic I, Doerks T, and Bork P. (2014). SMART: Recent updates, new developments and status in 2015. Nucleic Acids Research.43:D257–D260.

Linder JU. (2005). Substrate selection by class III adenylyl cyclases and guanylyl cyclases. IUBMB life.57:797–803.

Linder JU, Engel P, Reimer A, Krüger T, Plattner H, Schultz A, and Schultz JE. (1999). Guanylyl cyclases with the topology of mammalian adenylyl cyclases and an N□terminal P□type ATPase□like domain in Paramecium, Tetrahymena and Plasmodium. The EMBO Journal.18:4222–4232.

Linder JU, and Schultz JE. (2003). The class III adenylyl cyclases: Multi-purpose signalling modules. Cellular Signalling.15:1081–1089.

Long S, Brown KM, Drewry LL, Anthony B, Phan IQ, and Sibley LD. (2017). Calmodulin-like proteins localized to the conoid regulate motility and cell invasion by Toxoplasma gondii. PLoS Pathogens.13:e1006379.

Lourido S, Tang K, and Sibley LD. (2012). Distinct signalling pathways control Toxoplasma egress and host7cell invasion. The EMBO Journal.31:4524–4534.

Lucas KA, Pitari GM, Kazerounian S, Ruiz-Stewart I, Park J, Schulz S, Chepenik KP, and Waldman SA. (2000). Guanylyl cyclases and signaling by cyclic GMP. Pharmacological Reviews.52:375–414.

MacFarland RT. (1995). Molecular aspects of cyclic GMP signaling. Zoological Science.12:151–163.

Marchler-Bauer A, and Bryant SH. (2004). CD-Search: Protein domain annotations on the fly. Nucleic Acids Research.32:W327–W331.

Nitzsche R, Günay-Esiyok Ö, Tischer M, Zagoriy V, and Gupta N. (2017). A plant/fungal-type phosphoenolpyruvate carboxykinase located in the parasite mitochondrion ensures glucose-independent survival of Toxoplasma gondii. Journal of Biological Chemistry.292:15225–15239.

Pilz RB, and Casteel DE. (2003). Regulation of gene expression by cyclic GMP. Circulation Research.93:1034–1046.

Plattner F, Yarovinsky F, Romero S, Didry D, Carlier MF, Sher A, and Soldati-Favre D. (2008). Toxoplasma profilin is essential for host cell invasion and TLR11-dependent induction of an interleukin-12 response. Cell host & Microbe. Feb 14;3:77–87. Epub 2008/03/04.

Potter LR. (2011). Guanylyl cyclase structure, function and regulation. Cellular Signalling.23:1921–1926.

Roelofs J, Snippe H, Kleineidam RG, and Van Haastert P. (2001). Guanylate cyclase in Dictyostelium discoideum with the topology of mammalian adenylate cyclase. Biochemical Journal.354:697.

Rohloff P, Miranda K, Rodrigues JC, Fang J, Galizzi M, Plattner H, Hentschel J, and Moreno SN. (2011). Calcium uptake and proton transport by acidocalcisomes of Toxoplasma gondii. PLoS One.6:e18390.

Sidik SM, Hackett CG, Tran F, Westwood NJ, and Lourido S. (2014). Efficient genome engineering of Toxoplasma gondii using CRISPR/Cas9. PloS One.9:e100450.

Sievers F, Wilm A, Dineen D, Gibson TJ, Karplus K, Li W, Lopez R, McWilliam H, Remmert M, and Söding J. (20119. Fast, scalable generation of high□quality protein multiple sequence alignments using Clustal Omega. Molecular Systems Biology.7:539.

Sinha S, and Sprang S. (2006). Structures, mechanism, regulation and evolution of class III nucleotidyl cyclases. In: Reviews of Physiology Biochemistry and Pharmacology. Springer. p. 105–140.

Sonnhammer EL, Von Heijne G, and Krogh A. A hidden Markov model for predicting transmembrane helices in protein sequences. Proceedings of the Ismb; (1998).

Steegborn C. (2014). Structure, mechanism, and regulation of soluble adenylyl cyclases—Similarities and differences to transmembrane adenylyl cyclases. Biochimica Et Biophysica Acta (BBA)-Molecular Basis of Disease.1842:2535–2547.

Taylor HM, McRobert L, Grainger M, Sicard A, Dluzewski AR, Hopp CS, Holder AA, and Baker DA. (2010). The malaria parasite cyclic GMP-dependent protein kinase plays a central role in blood-stage schizogony. Eukaryotic Cell.9:37–45.

Tesmer JJ, Sunahara RK, Gilman AG, and Sprang SR. (1997). Crystal structure of the catalytic domains of adenylyl cyclase in a complex with Gsα• GTPγS. Science.278:1907–1916.

Yuasa K, Mi-Ichi F, Kobayashi T, Yamanouchi M, Kotera J, Kita K, and Omori K. (2005). PfPDE1, a novel cGMP-specific phosphodiesterase from the human malaria parasite Plasmodium falciparum. Biochemical Journal.392:221–229.

