## Supplementary material for "An unusual and physiologically vital protein with guanylate cyclase and P-type ATPase like domain in a pathogenic protist"

#### Supplementary information

**Figure S1. Phylogenetic analysis reveals a protozoan-specific clading of *Tg*ATPase$_P$-GC.** The phylogenetic tree shows the evolutionary relationship of *Tg*ATPase$_P$-GC from *T. gondii* with respective orthologs from various organisms representing the indicated domains of life. The image represents a single most parsimonious phylogenetic tree generated by UPGMA method. Sequence alignment and construction of the tree were performed by CLC Sequence Viewer 7.7 followed by visualization using Figtree v1.4.3 program. The colored dots on branching nodes show the bootstrap values for individual branches. Accession numbers: *Arabidopsis thaliana*, AAM51559.1; *Ascaris suum* (receptor-type), ERG86823.1; *Bos taurus* (retinal-type) NP_776973.2; *Bos taurus* (soluble α1-subunit), P19687.1; *Bos taurus* (soluble β1-subunit), P16068.1; *Caenorhabditis elegans* (receptor-type), NP_494995.2; *Caenorhabditis elegans* (soluble), NP_510557.3; *Drosophila melanogaster* (receptor-type), AAA85858.1; *Drosophila melanogaster* (soluble β-subunit), NP_524603.2; *Eimeria tenella* (soluble α), XP_013229212.1; *Eimeria tenella* (particulate β), XP_013235760.1; *Hammondia hammondi*, XP_008885058.1; *Homo sapiens* (retinal-type), NP_000171.1; *Homo sapiens* (soluble α1-subunit), NP_001124157.1; *Homo sapiens* (soluble β1-subunit), NP_001278880.1; *Musca domestica* (receptor-type), XP_005177218.1; *Musca domestica* (soluble), XP_019895151.1; *Mus musculus* (retinal-type), NP_001007577.1; *Mus musculus* (soluble α1-subunit), AAG17446.1; *Mus musculus* (soluble β1-subunit), AAG17447.1; *Oryza sativa*, ABD18448.1; *Oxytricha trifallax*, EJY85073.1; *Paramecium tetraurelia*, XP_001346995.1; *Plasmodium falciparum* (soluble α), AJ245435.1; *Plasmodium falciparum* (particulate β), AJ249165.1; *Prunus persica*, AGN29346.1; *Tetrahymena pyriformis*, AJ238858.1; *Toxoplasma gondii*, EPR59074.1; *Toxocara canis* (receptor-type), KHN81453.1; *Toxocara canis* (soluble), KHN85312.1.

**Figure S2. Sequence alignment of GC1 and GC2 domains of *Tg*ATPase$_P$-GC with other exemplary cyclases identifies signature residues**. Amino acid sequences were aligned with the Clustal Omega program. The color-coding refers to at least 30% sequence conservation. The secondary structures depicted underneath the alignment correspond to transmembrane type III adenylate cyclases (tmACs) (*Oryctolagus cuniculus* for GC1, *Rattus norvegicus* for GC2; DSSP analysis). In type III cyclases, 7 key residues are involved in cofactor binding for catalysis, which are indicated above the alignments as 'Me' for metal, 'B' for base, 'Pγ' for phosphate, 'R' for ribose, and 'Tr' for transition state binding. Note that alignment of GC1 and GC2 domains from *Tg*ATPase$_P$-GC to their orthologous GCs/ACs showed that unlike other cyclases, a 74-residues long segment (3033-3107 aa) is inserted between α3 and β4 of GC1 (not shown). Accession numbers: *Tg*ATPase$_P$-GC, *Toxoplasma gondii* (Uniprot S7VVK4); *Pf*GCα, *Plasmodium falciparum* (AJ245435.1); *Pf*GCβ, *P. falciparum* (AJ249165.1); *Pt*GC, *Paramecium tetraurelia* (XP_001346995.1; *Tp*GC, *Tetrahymena pyriformis* (AJ238858.1); soluble *Mm*GCα, *Mus musculus* (AAG17446.1); soluble *Mm*GCβ, *M. musculus* (AAG17447.1); retinal-type *Mm*GC2, *M. musculus* (NP_001007577.1); type II *Rn*tmAC, *Rattus norvegicus* (P26769.1) and; type V *Oc*tmAC, *Oryctolagus cuniculus* (CAA82562.1).

**Figure S3. Recombinant expression of *Tg*ATPase$_P$-GC1 and GC2 domains in M15 and BTH101 strains of *Escherichia coli.* (A)** Schematics depicting the molecular cloning of GC1, GC2 and GC1+GC2 domains in the *pQE60* expression vector. The open reading frames of indicated domains were amplified starting from the first upstream start codon (ATG) using tachyzoite-derived RNA and ligated into *Bgl*II-digested *pQE60* plasmid. Proteins were fused with a C-terminal 6xHis-tag for subsequent detection by immunoblot and purification by virtue of the Ni-NTA column. **(B)** Immunoblot of purified GC1-6xHis and GC2-6xHis proteins (5 μg) using α-His antibody. Purification was performed under denaturing conditions from the M15 strain, as described in *methods.* Purification of GC1+GC2 was not successful. The protein bands of 44-kDa and 34-kDa correspond to GC1 and GC2 domains, respectively. **(C)** Functional testing of GC1, GC2 and GC1+GC2 by complementation of the BTH101 strain were deficient in cAMP signaling. The transgenic strains expressing GC1, GC2, GC1+GC2, *Ec*CyaA (positive control), or harboring the empty *pQE60* vector (negative control) were grown in LB medium ($OD_{600}$,1.6; 30°C), and then dilution plated on MacConkey agar containing 1% maltose (carbon source) and 200 µM IPTG (inducer). The appearance of red colonies indicates a cAMP-dependent catabolism of disaccharide following a functional expression of *Ec*CyaA, but not others.

**Figure S4. Pharmacological inhibition of phosphodiesterases can repair phenotypic defects in the *Tg*ATPase$_P$-GC mutant**. **(A)** Gliding motility of the mutant ($P_{native}$-*Tg*ATPase$_P$-GC-HA$_{3'IT}$-3'UTR$_{excised}$) and progenitor ($P_{native}$-*Tg*ATPase$_P$-GC-HA$_{3'IT}$-3'UTR$_{floxed}$) strains in the presence of two inhibitors of cGMP-specific phosphodiesterases, zaprinast (500 µM) and BIPPO (55 µM). Fresh syringe-released parasites were treated with drugs for 15 min during the motility assay, followed by fixation and staining with α-*Tg*Sag1 antibody. 500-600 parasites of each strain were evaluated for the motile fraction, and 100-120 trail lengths were measured by ImageJ (n= 3 assays). **(B-C)** Egress and invasion rates of indicated parasite strains after exposure to zaprinast and BIPPO. Intracellular and extracellular parasites were differentially stained with α-*Tg*Gap45 and α-*Tg*Sag1 antibodies, as shown in *methods*. Drug treatment was performed during invasion assay for 1 h. For egress, parasitized cells (MOI, 1; 40 h post-infection) were stimulated with either zaprinast (500 µM) or BIPPO (55 µM) for 5:30 min prior to fixation and staining. In total, 1000 parasites and 500-600 vacuoles were scored for each strain in *panel b* and *c,* respectively (n= 3 assays). Numerical values in all graphs show the means with S.E.M. Statistics was done for individual pair of columns using Student's *t*-test (*, $p \leq 0.05$; **, $p \leq 0.01$; ***, $p \leq 0.001$).

**Figure S5**. **C-terminal epitope-tagging and Cre recombinase-mediated knockdown of *Tg*PKG in *T. gondii.*** **(A)** 3'-insertional tagging of the *Tg*PKG gene with a hemagglutinin tag (step 1) and subsequent deletion of loxP-flanked 3'UTR by Cre recombinase (step 2). The construct for 3'-insertional tagging (3'IT) *via* indicated crossover sequence (COS) was transformed into the parental strain (RHΔ*ku80-hxgprt*$^-$) and drug-selected for introduction of HXGPRT selection cassette (S.C.). The eventual strain (termed as progenitor, $P_{native}$-*Tg*PKG-HA$_{3'IT}$-3'UTR$_{floxed}$) expressed *Tg*PKG-HA$_{3'IT}$ under its own regulatory elements. In the second step, the progenitor strain was transfected with a vector expressing Cre-recombinase to excise the floxed 3'UTR and HXGPRT *via* negative selection, resulting in downregulation of *Tg*PKG expression. **(B)** Genomic screening PCR validating the integrity of the *Tg*PKG mutant generated by excision of 3'UTR. Primers indicated as red-color arrows in *panel a* were used to test gDNAs isolated from the mutant clones (C1-C3) and progenitor strain. A successful mutagenesis was further confirmed by sequencing of amplicons (1.9-kb). **(C)** Immunoblot showing the expression level of *Tg*PKG in two selected mutant clones along with the progenitor and parental strains. Extracellular tachyzoites ($2x10^7$) of each strain were subjected to protein isolation followed by immunoblotting with α-HA plus α-*Tg*Rop2 (loading control) antibodies. The presence of two *Tg*PKG isoforms (112-kDa and 135-kDa) in the progenitor, but not in the parental strain, confirms an efficient 3'-HA tagging (see *panel a*).

**Figure S6. Inhibition of residual activity in the *Tg*PKG-HA$_{3'IT}$ mutant by compound 2 aggravates the motility defect.** The motile fraction and the average trail length of the mutant ($P_{native}$-*Tg*PKG-HA$_{3'IT}$-3'UTR$_{excised}$) and progenitor ($P_{native}$-*Tg*PKG-HA$_{3'IT}$-3'UTR$_{floxed}$) strains, generated according to Figure S5, were measured by staining with α-*Tg*Sag1 antibody after compound 2 treatment (2 µM, 15 min). A total of 500 parasites for each strain were analyzed to calculate the motile fraction. In total, 100 trails were measured for the progenitor strain, whereas only 30 trails could be evaluated for the mutant due to severe defect. Numerical values show the means with S.E.M. Statistical significance was calculated for each pair of column individually by Student's *t*-test. (*, $p \leq 0.05$; **, $p \leq 0.01$; ***, $p \leq 0.001$).

**Table S1. Oligonucleotide sequences used in this study**

| Primer name (Restriction site) | Primer Sequence (Restriction site underlined) | Objective (Cloning vector) |
|---|---|---|
| *Tg*ATPase$_P$-GC-COS-F1 (*Xcm*I) | CTCATCCCACCGGTCACCTGGGC ATGAGTGTGGCGGAGT | Cloning of crossover sequence for 3'-HA tagging (*p3'IT-HXGPRT*) |
| *Tg*ATPase$_P$-GC-HA$_{3'IT}$ -COS-R1 (*EcoR*I) | CTCATCGAATTCCTACGCGTAGT CCGGGACGTCGTACGGGTACGAC CCGAGTGCAGAGC | |
| *Tg*PKG-COS-F1 (*Nco*I) | CTCATCCCATGGGAAAACTCGTT TTCCCGC | |
| *Tg*PKG-HA$_{3'IT}$-COS-R1 (*EcoR*I) | CTCATCGAATTCCTACGCGTAGT CCGGGACGTCGTACGGGTAGAAA TCCTTGTCCCAGTCATACT | |
| *Tg*ATPase$_P$-GC-3`UTR-F1 (*EcoR*I+loxP) | CTCATCGAATTCATAACTTCGTAT AGCATACATTATACGAAGTTATA ACGCAGCTTTTGTCAGC | Cloning of 3'UTR for the native expression (*p3'IT-HXGPRT)* |
| *Tg*ATPase$_P$-GC-3`UTR-R1 (*Spe*I) | CTCATCACTAGTTTATGTACGTAT ATACGCACATGTATG | |
| *Tg*PKG-3`UTR-F1 (*EcoR*I+loxP) | CTCATCGAATTCATAACTTCGTAT AGCATACATTATACGAAGTTATT TTTTCAGCTTAGGTGTTTGTTCC | |
| *Tg*PKG-3`UTR-R1 (*Spe*I) | CTCATCACTAGTGCTTTTCTGCG ACTCTGCTC | |
| *Tg*ATPase$_P$-GC-HA$_{3'IT}$-Scr-F1 | CTGGTCTCCGCAGAGATGCT | Screening primers of 3'UTR excision for the gene knockdown *(p3'IT-HXGPRT)* |
| Scr-*Tg*PKG-HA$_{3'IT}$-Scr-F1 | GTTCATGTGCGGACCTCTCC | |
| 3`UTR$_{excised}$-loxP-Scr-R1 (*Tg*ATPase$_P$-GC/*Tg*PKG) | CAGTGAGCGCAACGCAATTA | |
| *Tg*ATPase$_P$-GC-CyCc1-F1 (*Bgl*II) | CTCATCAGATCTATGCTCGATAA GAAGTACTTGCCCCCAC | Cloning of *Tg*ATPase$_P$-GC cyclase catalytic domains for *E. coli* expression (*p*QE60-6xHis) |
| *Tg*ATPase$_P$-GC-CyCc1-R1 (*Bgl*II) | CTCATCAGATCTCGACGACGCAC CCGCAGT | |
| *Tg*ATPase$_P$-GC-CyCc2-F1 (*Bgl*II) | CTCATCAGATCTATGACGATGAG CTTAACGTTCATCATC | |
| *Tg*ATPase$_P$-GC-CyCc2-R1 (*Bgl*II) | CTCATCAGATCTTTGAGGGATCG CACCGCC | |
| *Tg*ATPase$_P$-GC-CyCc1+2-F1 (*Bgl*II) | CTCATCAGATCTATGCTCGATAA GAAGTACTTGCCCCCAC | |
| *Tg*ATPase$_P$-GC-CyCc1+2-R1 (*Bgl*II) | CTCATCAGATCTTTGAGGGATCG CACCGCC | |
| *Tg*ATPase$_P$-GC-KO-5'UTR-F1 (*Xcm*I) | CTCATCCCACCGGTCACCTGGTG CTTGGCTGATTGATGG | Cloning of 5'UTR for the *Tg*ATPase$_P$-GC knockout (*pKO-DHFR-TS*) |
| *Tg*ATPase$_P$-GC-KO-5'UTR-R1 (*Spe*I) | CTCATCACTAGTTTTCGTCGTATT CGATAGCTCC | |
| *Tg*ATPase$_P$-GC-KO-5'Scr-F1 | CGAGACGAGATTCTTGAACGA | Screening of 5' recombination |

| | | |
|---|---|---|
| *Tg*ATPase$_P$-GC-KO-3'UTR-F1 (*Hind*III) | CTCATCAAGCTTAACGCAGCTTTTGTCAGCG | Cloning of 3'UTR for the *Tg*ATPase$_P$-GC knockout (*pKO-DHFR-TS*) |
| *Tg*ATPase$_P$-GC-KO-3'UTR-R1 (*Apa*I) | CTCATCGGGCCCGACGGCTAGTCGAGACCCTG | |
| *Tg*ATPase$_P$-GC-KO-3'Scr-R1 | AAGGATGATATTGCACCTCACA | Screening of 3' recombination |
| *Tg*ATPase$_P$-GC-sgRNA-F1 | AAGTTCGTTGACTCTGTTCACCGCCG | Cloning of sgRNA- for editing of *Tg*ATPase$_P$-GC (*pU6-sgRNA-Cas9*) |
| *Tg*ATPase$_P$-GC-sgRNA-R1 | AAAACGGCGGTGAACAGAGTCAACGA | |

### Figure S1

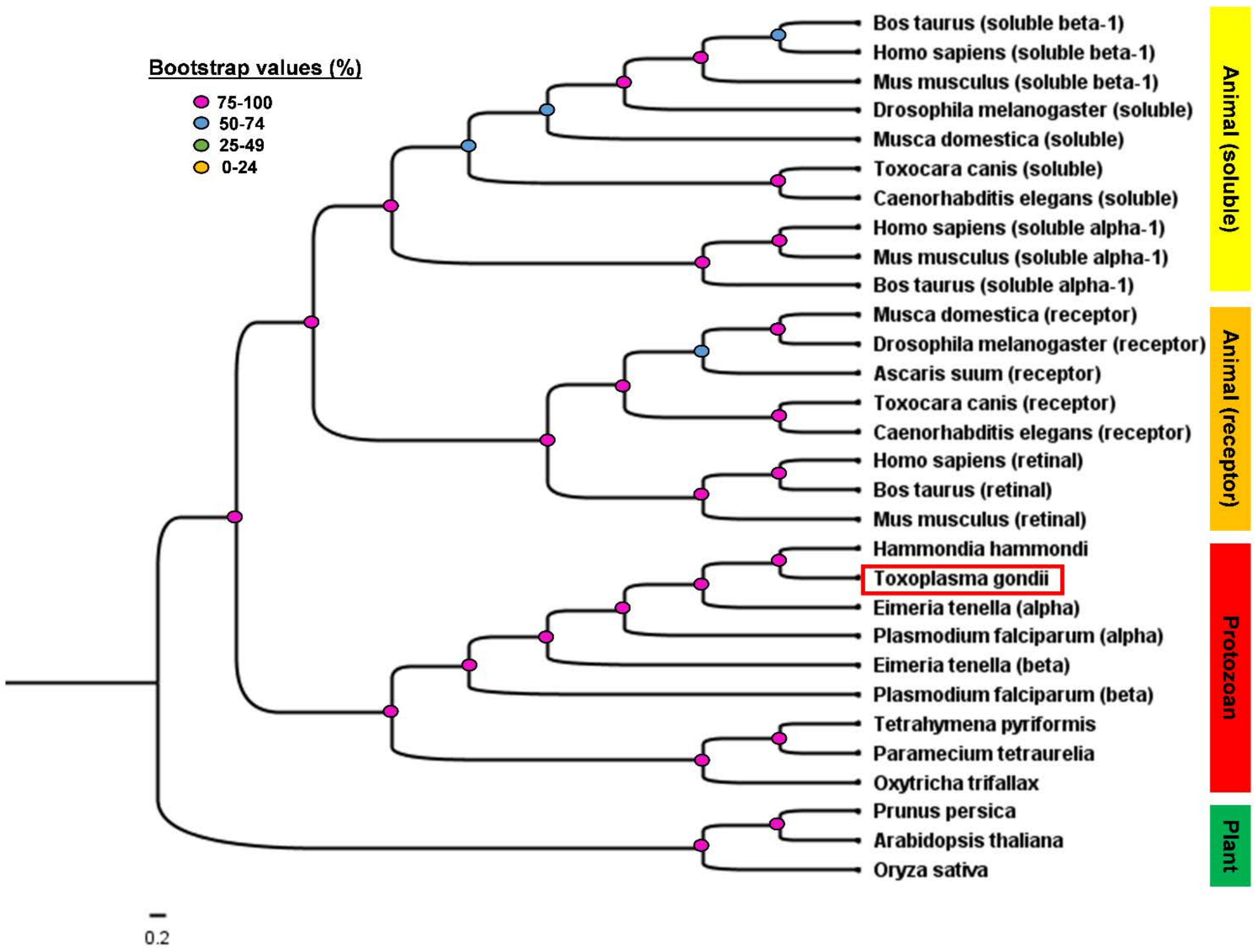
Bootstrap values (%)
75-100
50-74
25-49
0-24
Bos taurus (soluble beta-1)
Homo sapiens (soluble beta-1)
Mus musculus (soluble beta-1)
Drosophila melanogaster (soluble)
Musca domestica (soluble)
Toxocara canis (soluble)
Caenorhabditis elegans (soluble)
Homo sapiens (soluble alpha-1)
Mus musculus (soluble alpha-1)
Bos taurus (soluble alpha-1)
Musca domestica (receptor)
Drosophila melanogaster (receptor)
Ascaris suum (receptor)
Toxocara canis (receptor)
Caenorhabditis elegans (receptor)
Homo sapiens (retinal)
Bos taurus (retinal)
Mus musculus (retinal)
Hammondia hammondi
Toxoplasma gondii
Eimeria tenella (alpha)
Plasmodium falciparum (alpha)
Eimeria tenella (beta)
Plasmodium falciparum (beta)
Tetrahymena pyriformis
Paramecium tetraurelia
Oxytricha trifallax
Prunus persica
Arabidopsis thaliana
Oryza sativa
Animal (soluble)
Animal (receptor)
Protozoan
Plant
0.2

Figure S2

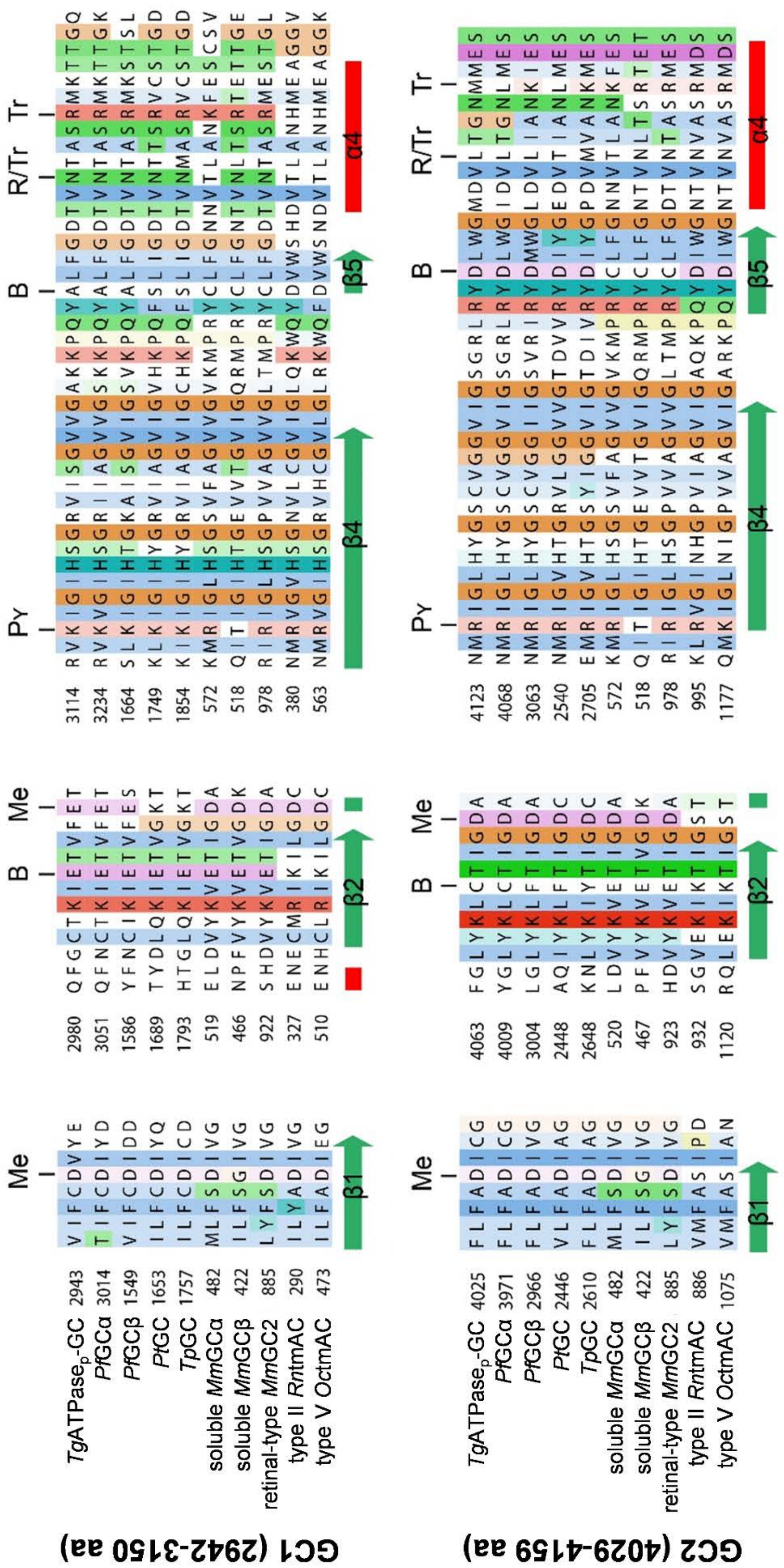
GC1 (2942-3150 aa)
GC2 (4029-4159 aa)
TgATPasep-GC
PfGCα
PfGCβ
PtGC
TpGC
soluble MmGCα
soluble MmGCβ
retinal-type MmGC2
type II RnAC
type V OctmAC
Me
B
Pγ
R/Tr
Tr
β1
β2
β4
β5
α4

### Figure S3

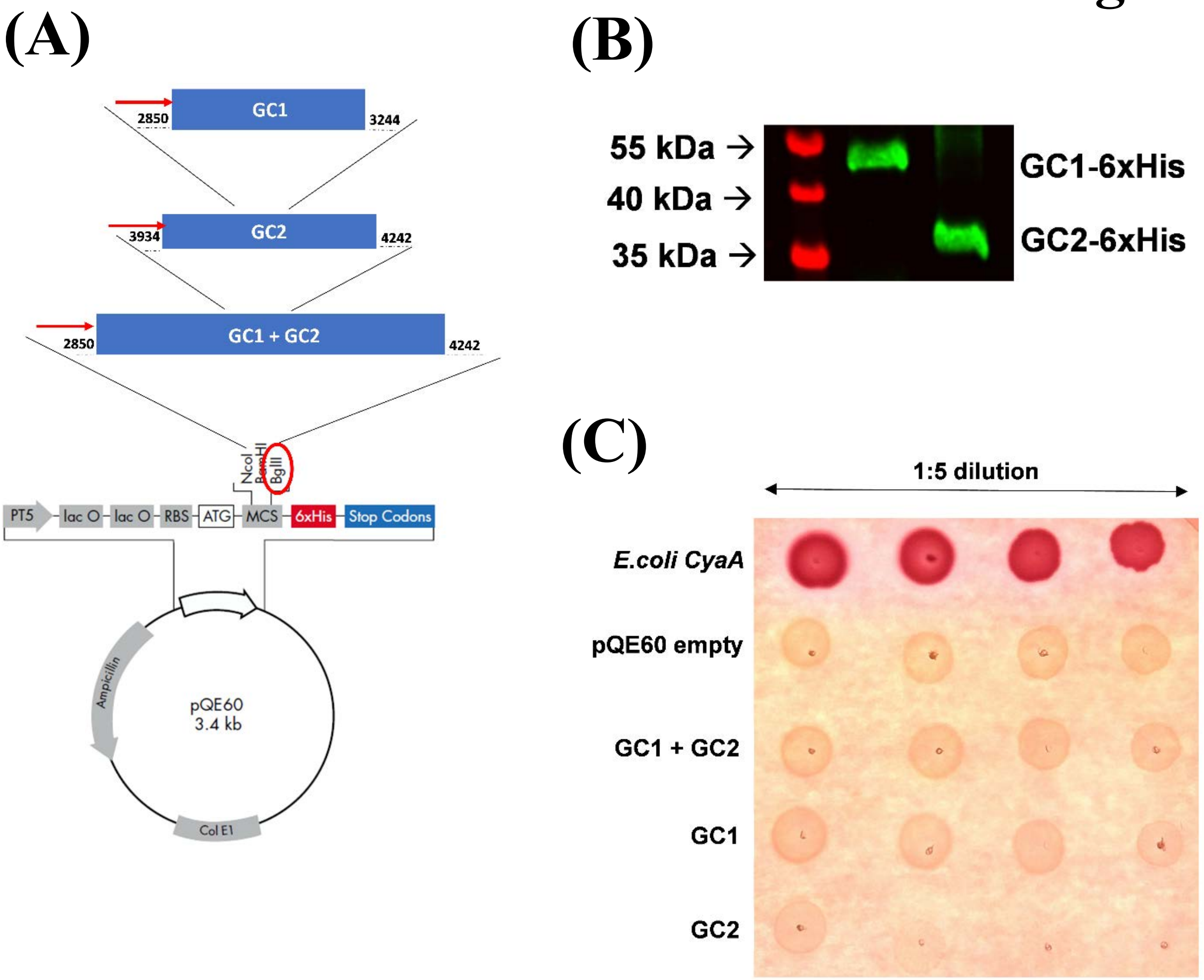
(A)
GC1
2850
3244
GC2
3934
4242
GC1 + GC2
2850
4242
NcoI
BamHI
BglII
PT5
lac O
lac O
RBS
ATG
MCS
6xHis
Stop Codons
Ampicillin
pQE60
3.4 kb
Col E1
(B)
55 kDa →
40 kDa →
35 kDa →
GC1-6xHis
GC2-6xHis
(C)
1:5 dilution
E.coli CyaA
pQE60 empty
GC1 + GC2
GC1
GC2

### Figure S4

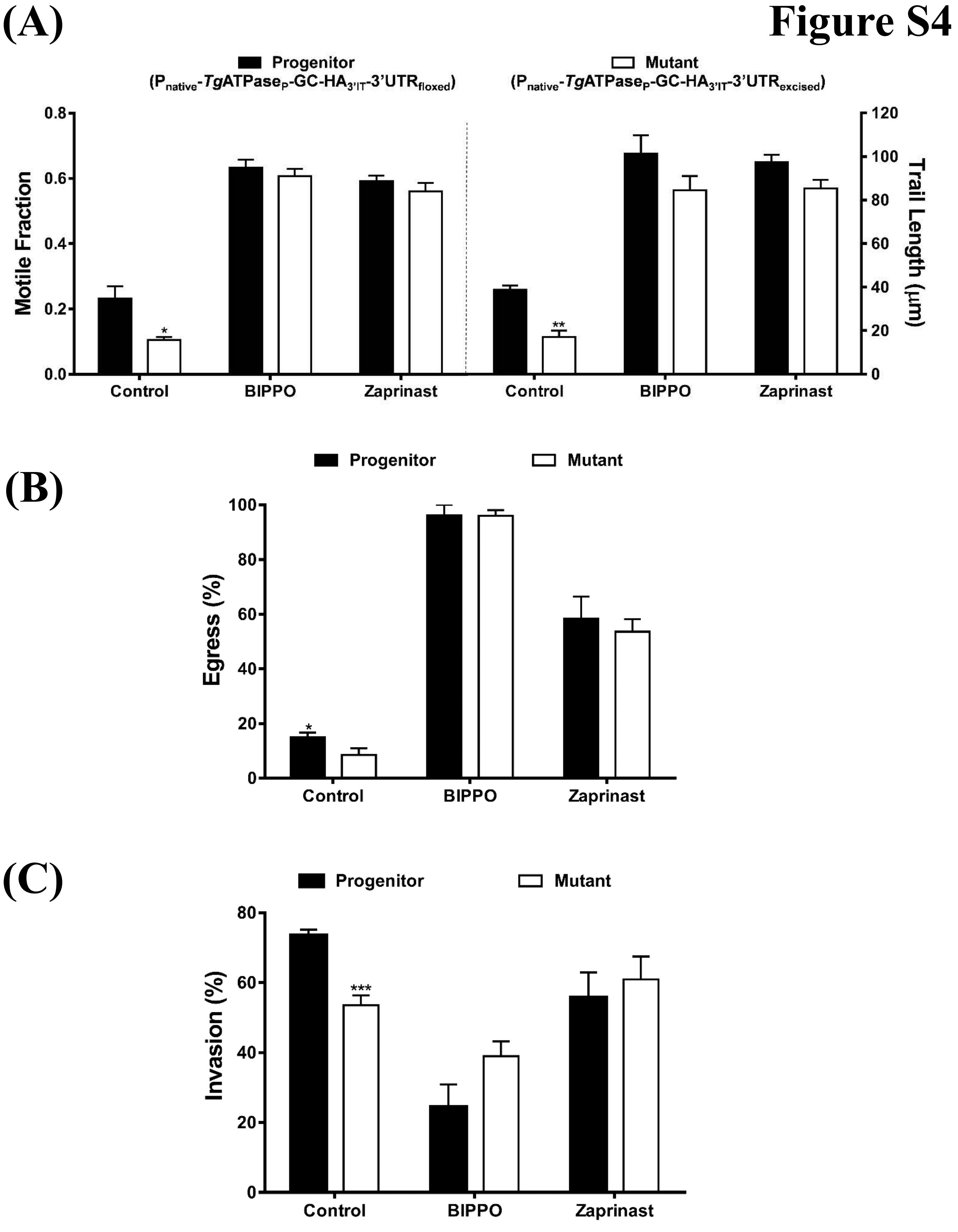
(A)
Progenitor
(Pnative-TgATPaseP-GC-HA3'IT-3'UTRfloxed)
Mutant
(Pnative-TgATPaseP-GC-HA3'IT-3'UTRexcised)
Motile Fraction
Trail Length (μm)
Control
BIPPO
Zaprinast
(B)
Progenitor
Mutant
Egress (%)
Control
BIPPO
Zaprinast
(C)
Progenitor
Mutant
Invasion (%)
Control
BIPPO
Zaprinast

### Figure S5

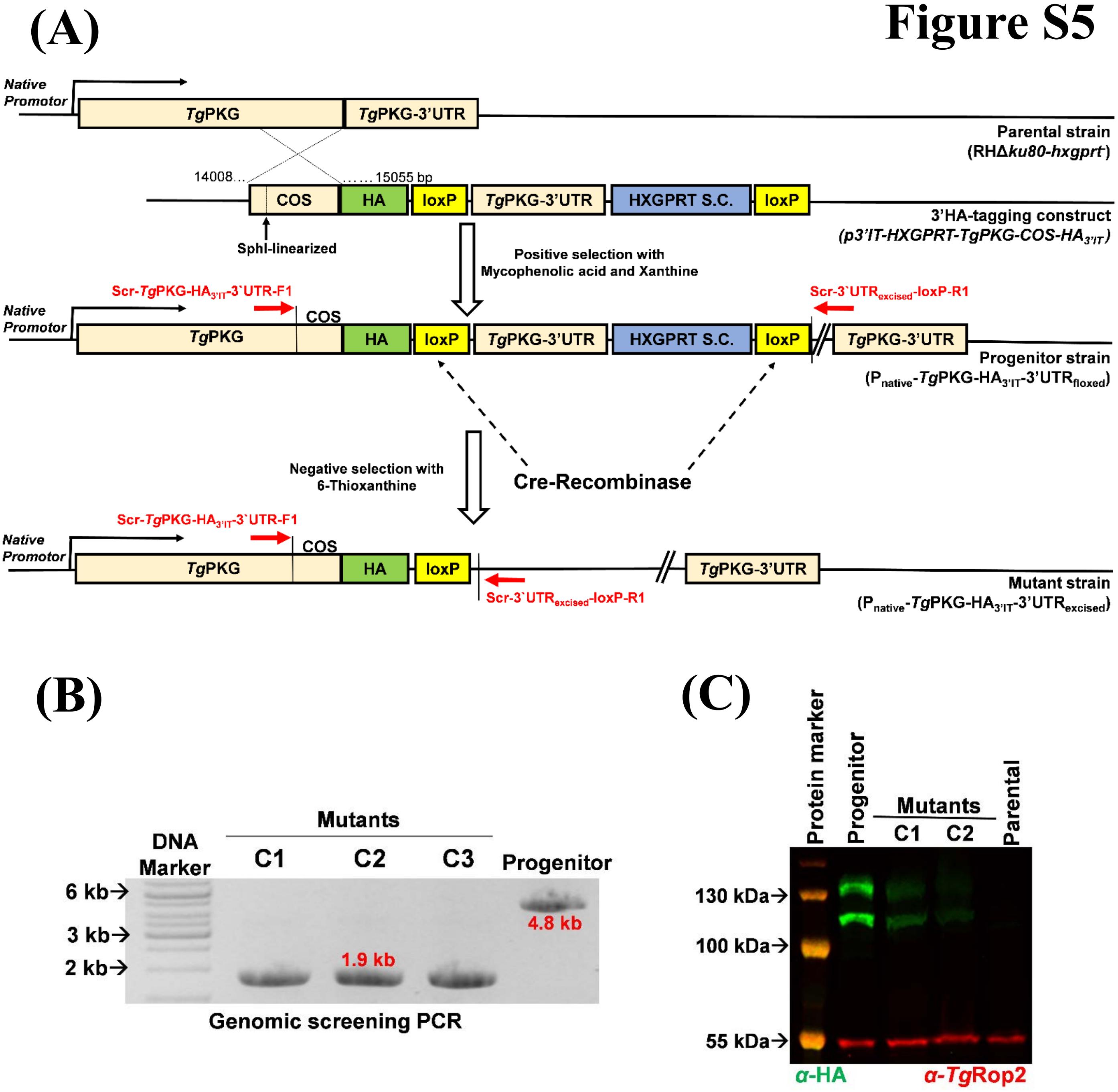
(A)
Native Promotor
TgPKG
TgPKG-3'UTR
Parental strain
(RHΔku80-hxgprt-)
14008…
……15055 bp
COS
HA
loxP
TgPKG-3'UTR
HXGPRT S.C.
loxP
3'HA-tagging construct
(p3'IT-HXGPRT-TgPKG-COS-HA3'IT)
SphI-linearized
Positive selection with Mycophenolic acid and Xanthine
Scr-TgPKG-HA3'IT-3`UTR-F1
Scr-3`UTRexcised-loxP-R1
Native Promotor
COS
TgPKG
HA
loxP
TgPKG-3'UTR
HXGPRT S.C.
loxP
TgPKG-3'UTR
Progenitor strain
(Pnative-TgPKG-HA3'IT-3'UTRfloxed)
Negative selection with 6-Thioxanthine
Cre-Recombinase
Scr-TgPKG-HA3'IT-3`UTR-F1
Native Promotor
COS
TgPKG
HA
loxP
TgPKG-3'UTR
Scr-3`UTRexcised-loxP-R1
Mutant strain
(Pnative-TgPKG-HA3'IT-3'UTRexcised)
(B)
Mutants
DNA Marker
C1
C2
C3
Progenitor
6 kb
3 kb
2 kb
1.9 kb
4.8 kb
Genomic screening PCR
(C)
Protein marker
Progenitor
Mutants
C1
C2
Parental
130 kDa
100 kDa
55 kDa
α-HA
α-TgRop2

### Figure S6

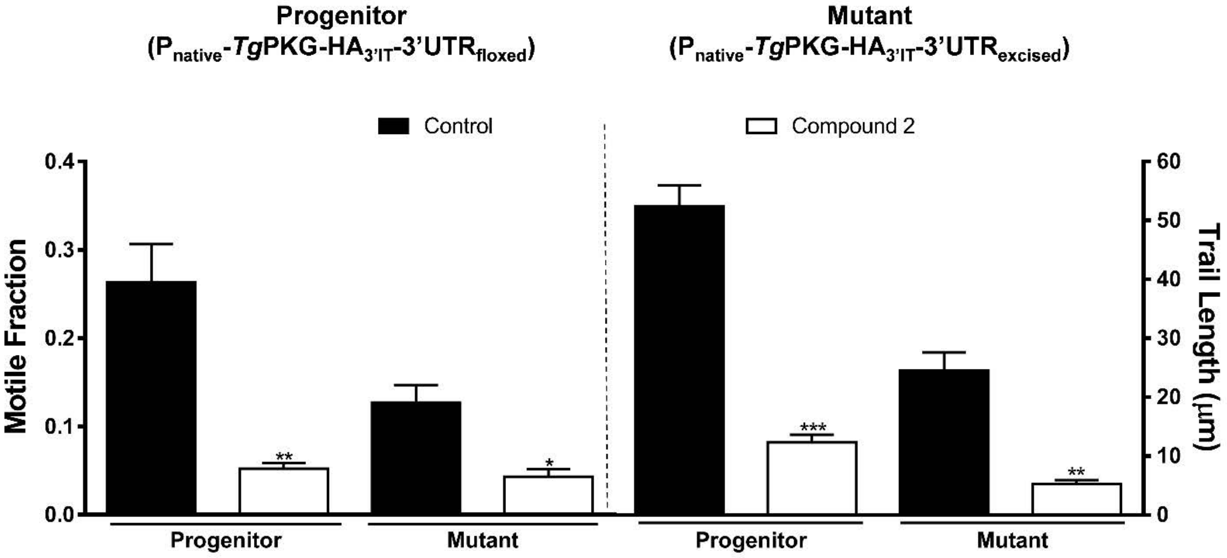
Progenitor
($P_{native}$-*Tg*PKG-$HA_{3'IT}$-3'$UTR_{floxed}$)
Mutant
($P_{native}$-*Tg*PKG-$HA_{3'IT}$-3'$UTR_{excised}$)
Control
Compound 2
Motile Fraction
0.4
0.3
0.2
0.1
0.0
Trail Length (μm)
60
50
40
30
20
10
0
**
*
***
**
Progenitor
Mutant
Progenitor
Mutant
